# Astrocytes close the critical period for visual plasticity

**DOI:** 10.1101/2020.09.30.321497

**Authors:** Jérôme Ribot, Rachel Breton, Charles-Félix Calvo, Julien Moulard, Pascal Ezan, Jonathan Zapata, Kevin Samama, Alexis-Pierre Bemelmans, Valentin Sabatet, Florent Dingli, Damarys Loew, Chantal Milleret, Pierre Billuart, Glenn Dallérac, Nathalie Rouach

## Abstract

Brain postnatal development is characterized by critical periods of experience-dependent remodeling^1,2^. Termination of these periods of intense plasticity is associated with settling of neuronal circuits, allowing for efficient information processing^3^. Failure to end critical periods thus results in neurodevelopmental disorders^4,5^. Yet, the cellular processes defining the timing of these developmental periods remain unclear. Here we show in the mouse visual cortex that astrocytes control the closure of the critical period. We uncover a novel underlying pathway involving regulation of the extracellular matrix that allows interneurons maturation via an unconventional astroglial connexin signaling. We find that timing of the critical period closure is controlled by a marked developmental upregulation of the astroglial protein connexin 30 that inhibits expression of the matrix degrading enzyme MMP9 through the RhoA-GTPase signaling pathway. Our results thus demonstrate that astrocytes not only influence activity and plasticity of single synapses, but are also key elements in the experience-dependent wiring of brain developing circuits. This work, by revealing that astrocytes promote the maturation of inhibitory circuits, hence provide a new cellular target to alleviate malfunctions associated to impaired closure of critical periods.

## Main text

During the first weeks of life, massive synaptogenesis occurs and is followed by shaping of synaptic circuits^6^. In the last decades, the role of astroglial processes as structural and signaling partners of individual synapses has been established and their implication in neuronal network activities and cognitive functions has recently been unveiled^7^. Yet, whether they take part in the wiring of the neuronal circuitry that occurs during critical periods of postnatal development remains unknown. The visual cortex is a hallmark brain region of experience-dependent shaping of synaptic circuits during a period of enhanced plasticity that follows eyes opening^2,8^. Intriguingly, one pioneer study published 30 years ago showed that introducing immature astrocytes in adult cats re-opens a period of high plasticity, reminiscent of the critical period^9^. However, since then, whether and how maturation of astrocytes actually take part in the control of the critical periods closure has never been investigated.

## Immature astrocytes favor plasticity

We first investigated the ability of immature astrocytes to promote visual cortex plasticity in adult mice. To this end, we cultured and labeled (lentivirus PGK-GFP) immature astrocytes from the cortex of P1-P3 (postnatal day 1-3) mice, and implanted them ten days later in the primary visual cortex (V1) of adult mice (P100), in which the critical period is closed (Fig. 1a). Two weeks after the graft, we tested mice for ocular dominance (OD) plasticity, a form of plasticity typical of the critical period where changes in visual inputs alter the natural dominance of the contralateral eye. To do so, we assessed visual cortex activity using optical imaging of intrinsic signals after four days of monocular deprivation (MD) (Fig. 1a). We found that mice engrafted with immature astrocytes displayed a high level of plasticity, unlike control mice subjected to MD with injection of culture medium or non-injected mice (Fig. 1b-c). These data indicate that immature astrocytes re-open OD plasticity in adult mice.

**Figure 1.**
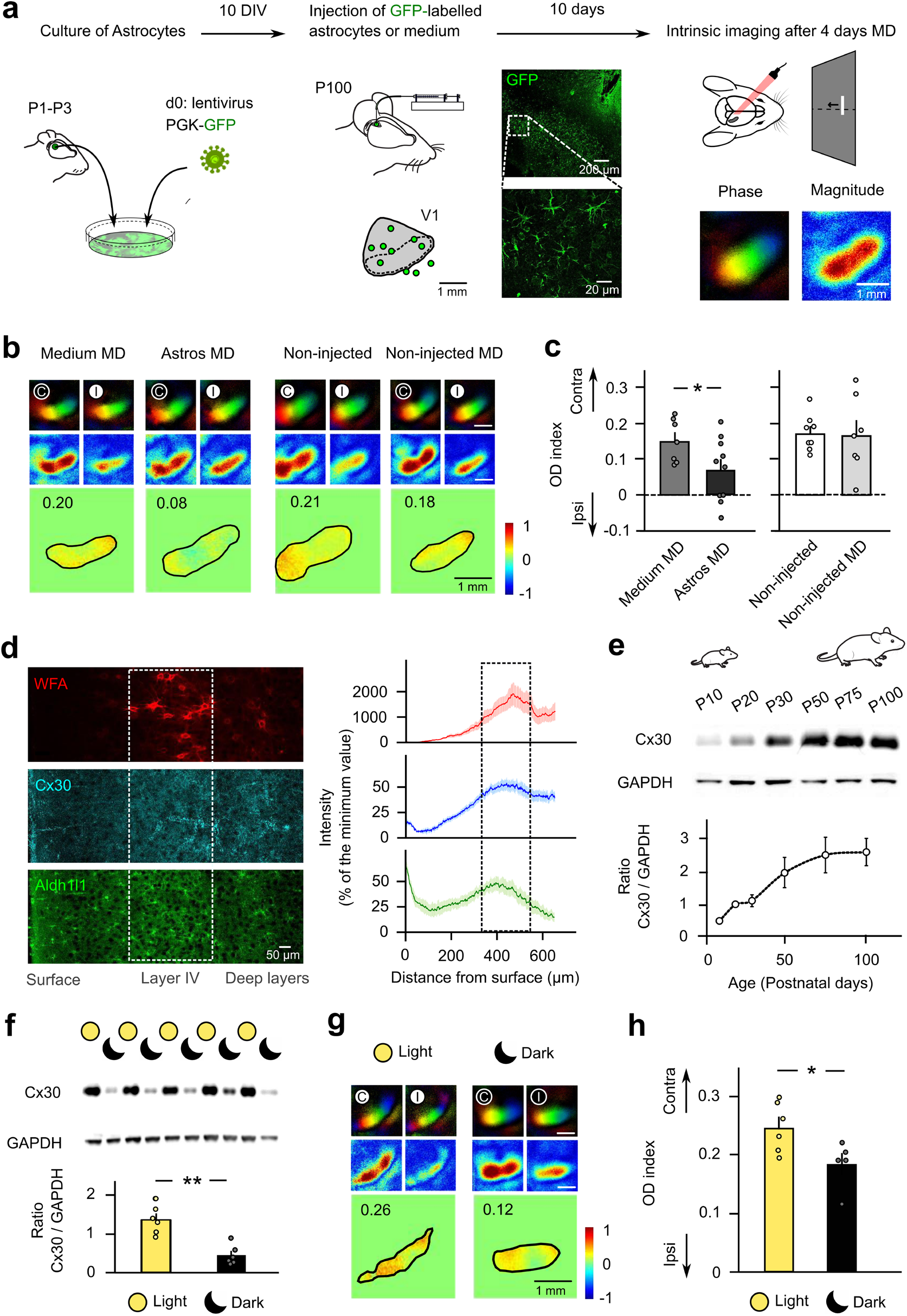
Immature astrocytes favor cortical plasticity. (a) Immature astrocytes from P1 - P3 mice were transduced with PGK-GFP lentivirus, cultured for 10 DIV, and injected in the V1 area of adult mice (P100), as shown on the schematic (green circles), where the dashed line represents the binocular zone. Confocal images of GFP-expressing astrocytes integrated in the visual cortex of injected mice are shown. Intrinsic optical imaging was then conducted to record phase and magnitude retinotopic maps for each eye stimulation after 4 days of monocular deprivation (MD). (b) These maps are shown in top panels for animals injected with control medium (Medium MD, n=7) or immature astrocytes (Astro MD, n=10) and for non-injected mice in control (non-injected, n=8) or MD (non-injected MD, n=7) conditions. Stimulated contralateral (C) and ipsilateral (I) eyes are indicated with a white (open) or black (closed) circle. Bottom panels show the normalized OD map with average value inset. (c) Mice injected with immature astrocytes showed a marked increase in OD plasticity compared with control mice injected with culture medium (P=0.043, DF=15, two-tailed t-test), while no effect was observed in non-injected mice (P=0.899, DF=13, two-tailed t-test). (d) Representative immunostaining for WFA (top), Cx30 (middle) and Aldh1l1 (bottom) in V1 of P50 mouse (left panels), and corresponding quantification of spatial distributions across V1 layers (right panels, n=3 mice). (e) Western blot analysis reveals changes in Cx30 levels in the visual cortex during development (n=6 per age, P=0.0001, Kruskal-Wallis (KW=22.31)) and (f) after 4 days of dark exposure (n=6 per group, P=0.0022, U=0, Mann-Whitney test). (g) Intrinsic optical imaging for light (n=6) and dark (n=5) exposed animals showing that (h) dark exposure decreased OD index (P=0.035, DF=9, two-tailed t-test).

To identify how immature astrocytes favor OD plasticity, we then investigated the molecular determinants of astroglial maturation. Comparing gene expression of immature (P7) *vs* mature (P30) astrocytes using the transcriptome database for astrocytes during development^10^ revealed about 200 genes that were differentially expressed with a fold-change over 5 (Extended data Table 1). Gene Ontology analysis of enriched astroglial gene groups identified a functional switch during maturation from cell division to cell communication, with the cell junction genes being the most represented (Extended Table 2 and Fig. 1). Among these genes, *Gjb6*, encoding the astroglial gap-junction channel subunit connexin 30 (Cx30), displayed one of the highest increase in expression (Fold-change=9, *P*=0.0274; Extended Data Table 1). Accordingly, we recently found that Cx30 regulates the structural maturation of hippocampal astrocytes during postnatal development^11^. Together, these data incited us to investigate the role of Cx30 in the astroglial control of the critical period. To do so, we first assessed the regional and temporal expression of Cx30 in the mouse V1. We found by immunohistochemistry that Cx30 is enriched in layer 4, the main V1 input layer, identified by Wisteria Floribunda Agglutinin (WFA), a marker of perineuronal nets important for the timecourse of the critical period^12^. This enrichment correlates with a high density of astrocytes labeled in the Aldh1l1-gfp mice (Fig. 1d). Moreover, Cx30 protein levels increased from P10 to P50, as shown by western blot (Fig. 1e), reaching its maximum when the critical period ends, thus suggesting that it may contribute to its closure. If so, re-opening a phase of high plasticity should be associated with a downregulation of Cx30. As a period of dark exposure (DE) can reinstate visual cortex plasticity during adulthood after closure of the critical period^13,14^, we placed wild-type adult mice (P50) in the dark for 4 days and quantified V1 Cx30 levels. We found that this manipulation drastically reduced Cx30 protein levels (~-70%) (Fig. 1f). Remarkably, we also found that the same DE for 4 days induced by itself V1 plasticity, as indicated by the change in OD index, which suggests that astroglial Cx30 is a brake to plasticity (Fig. 1g-h).

## Astroglial Cx30 closes the critical period

We then directly investigated whether Cx30 inhibits OD plasticity. To do so, we first generated an astroglial knockdown mouse line for Cx30 (KD), where Cx30 expression is decreased in V1 astrocytes by ~70% (Extended data Fig. 2). In these mice, electroretinograms were unaltered, suggesting normal retinal functions (Extended Fig. 3). We found that OD plasticity peaked at P28 in wild-type (WT) mice. In contrast, this plasticity continued to increase in KD mice until P50, indicating an impairment in the closure of the critical period (Fig. 2a, b). While in WT mice this plasticity was due to an increase in the open eye response, in KD mice, it resulted from a reduction of the closed eye response, which typically reflects changes occurring in mature and immature system, respectively^15^. These data thus indicate that Cx30 is required for proper maturation of the visual cortex.

**Figure 2.**
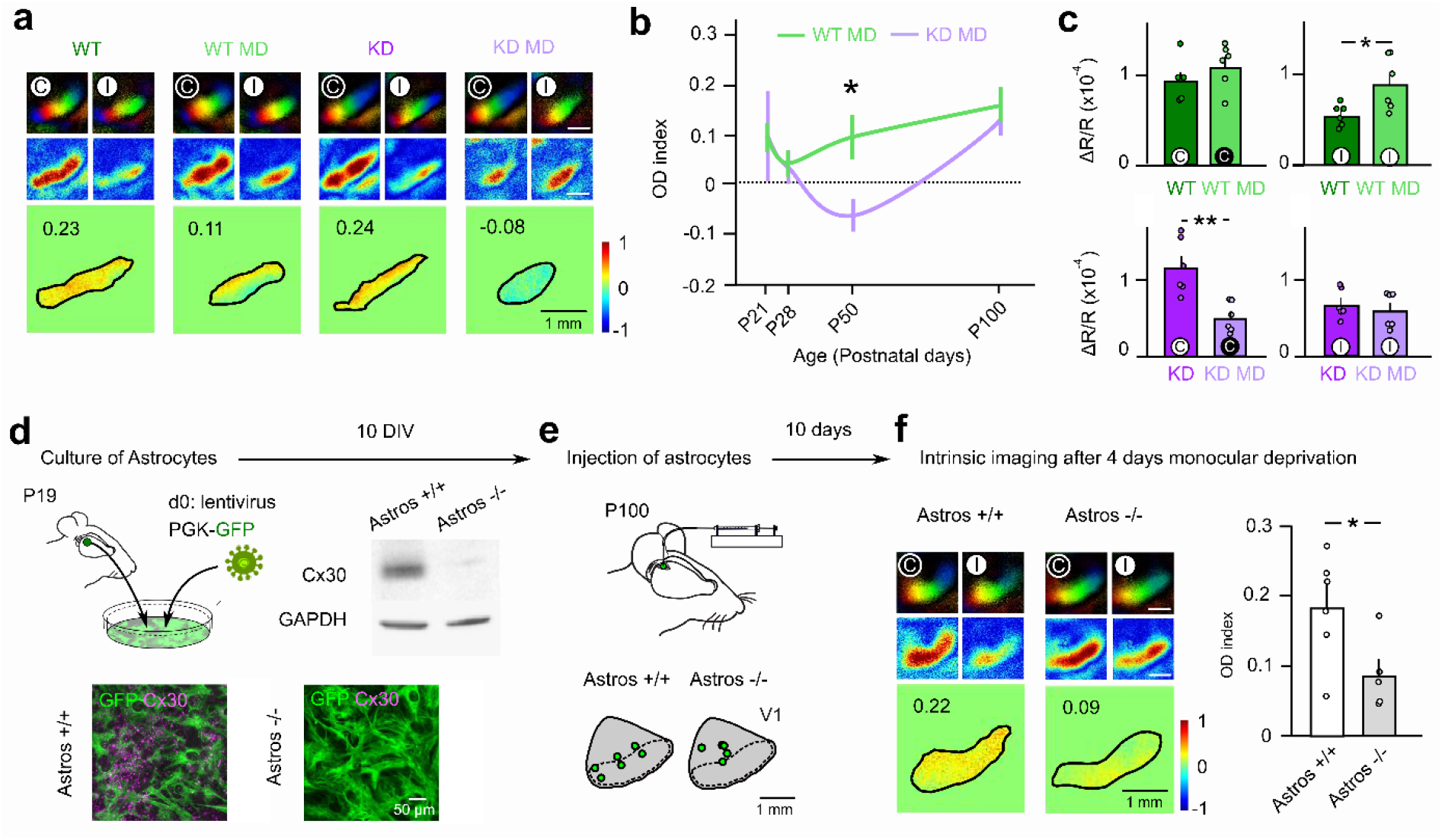
Astroglial connexin 30 closes the critical period. (a) Functional maps in P50 wild-type (WT) and Cx30 knockdown (KD) mice in control and MD conditions. (b) Developmental profile of OD index after MD in WT (P21, n=6; P28, n=7; P50, n=6; P100, n=7) and KD mice (P21, n=3; P28, n=8; P50, n=6; P100, n=11), indicating a protracted and enhanced plasticity at P50 in KD mice (P=0.0232, DF=46, two-way ANOVA followed by Sidak post-hoc test). (c) Changes in response magnitude after MD in P50 mice result from an increased response to the open (ipsilateral, I) eye stimulation in WT mice (n=6, P=0.0411, U=5, Mann Whitney test), and from a decreased response to the closed (contralateral, C) eye stimulation in KD mice (n=6, P=0.0022, U=0, Mann-Whitney test). (d) Mature astrocytes expressing or not Cx30 were isolated from P19 WT (Astro +/+) or constitutive Cx30 knockout mice (Astros −/−), respectively, and were then transduced with PGK-GFP lentiviruses and cultured for 10 DIV. Western blots and immunostaining show expression of Cx30 in Astro +/+, but not in Astro −/−. (e) The cultured astrocytes were injected in adult mice (P100), as shown on the schematics (green circles). (f) Functional maps show that OD plasticity is enhanced in mice injected with Astro −/− (n=5) compared to mice injected with Astro +/+ (n=6, P=0.038, DF=9, two-tailed t-test).

To further directly test that the expression of Cx30 in mature astrocytes inhibits OD plasticity, we engrafted mature astrocytes expressing or not Cx30, isolated from P19 wild type or constitutive Cx30 knockout mice, respectively (Fig. 2d, e). By doing so, we found that only the graft of mature astrocytes lacking Cx30 re-opened OD plasticity in adult mice (Fig. 2f). Altogether, these data indicate that astroglial Cx30 controls the timing of the critical period closure.

## Astroglial Cx30 promotes the maturation of inhibitory circuits

The temporal course of visual cortex critical period is determined by the maturation of local inhibitory circuits controlling the excitation-inhibition (E-I) balance^16-18^. To get insights on the physiological processes via which Cx30 closes the critical period, we thus measured changes in excitatory and inhibitory synaptic transmission following MD in pyramidal neurons from layer 4 visual cortex of WT and KD adult mice (P50) (Fig. 3a). The frequency, but not the amplitude, of both spontaneous excitatory and inhibitory postsynaptic currents (sEPSCs and sIPSCs) were markedly reduced in KD mice compared to WT (Fig. 3b-c). In addition, MD induced an increase in the frequency of sEPSC and sIPSCs in KD mice while it had no effect in WT, indicating that experience-dependent plasticity of both excitatory and inhibitory synapses occurs in adult KD, but not in WT mice. As excitation and inhibition influence each other through homeostatic processes^19^, we then assessed whether the E-I balance was affected by analyzing the dynamic conductances of evoked composite synaptic responses. We found the inhibition/excitation ratio to be reduced in KD mice (Fig. 3d-e), thus indicating functionally that inhibition is primarily affected by Cx30 downregulation. Further, as found for the sEPSC/sIPSC data, I-E ratio increased when plasticity was induced through MD. Together, these data suggest that experience-dependent maturation of inhibitory circuits ending the critical period requires astroglial Cx30. To investigate this possibility, we assessed maturity of PV interneurons, which settle visual cortex inhibition, by determining the abundance of their perineuronal nets (PNN) of extracellular matrix (ECM) proteins^12^. We found PNN to be significantly smaller in KD mice (Fig. 3f-g), thus revealing the immaturity of these cells. In addition, while MD also decreased PNN in WT mice, it had no further effect in KD mice, suggesting that reduction of PNN is a prerequisite for plasticity. Together, these data indicate that the developmental rise of Cx30 during the critical period is required for the timely maturation of visual cortex inhibition.

**Figure 3.**
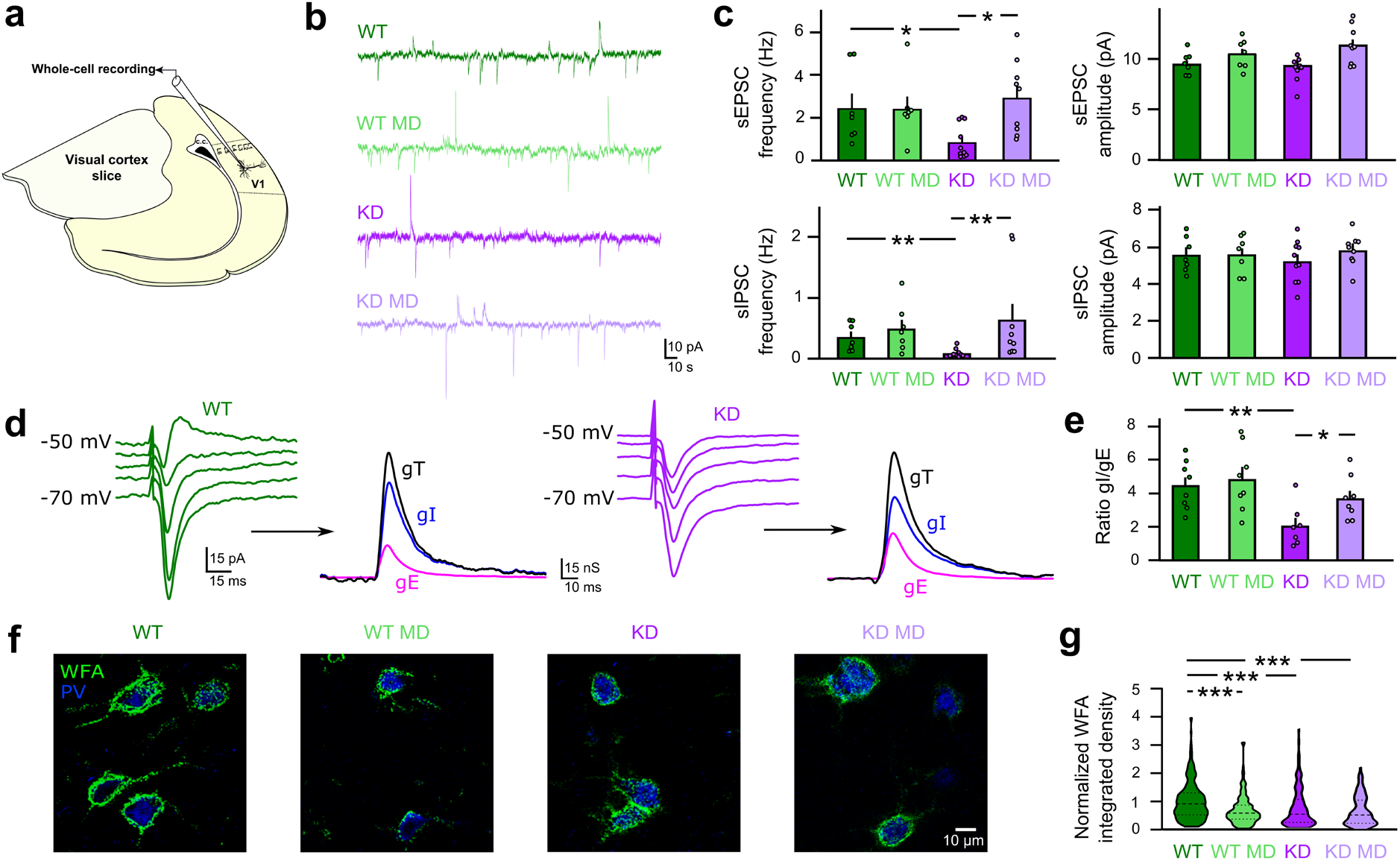
Astroglial Cx30 promotes the maturation of inhibitory circuits. (a) Schematic diagram depicting the brain slice from which V1 pyramidal neurons from layer 4 were recorded. (b) Example traces of sEPSCs (inward) and sIPSCs (outward) currents. (c) Frequency, but not amplitude, of both sEPSCs (P=0.0192, U=11, Mann-Whitney test) and sIPSCs (P=0.0034, U=5, Mann-Whitney test) was decreased in KD (n=10) compared to WT mice (n=7). MD induced no plasticity in WT mice (n=7, P=0.7027, U=19 for sEPSCs, P=0.6547 U=21, for sIPSCs, Mann-Whitney test), while it potentiated both the sEPSCs (P=0.0090, U=13, Mann-Whitney test) and sIPSCs (P=0.0019, U=13, Mann-Whitney test) in KD mice (n=9). (d) Example of evoked composite responses at increasing holding potentials allowing decomposition of the total conductance (gT) into excitatory (gE) and inhibitory (gI) conductances in WT and KD mice. (e) Inhibition/Excitation (I/E) ratio was reduced in KD mice (n=7, P=0.0078, U=5, Mann-Whitney test) compared to WT (n=8). MD induced no change in WT mice (n=8, P=0.7527, U=29, Mann-Whitney test), while it increased the I/E ratio in KD mice (n=8, P=0.020, U=8, Mann-Whitney test). (f) WFA immunostaining and (g) violin plot showing smaller perineuronal nets around PV interneurons in KD (n=177 cells from 16 mice, P<0.0001), WT MD (n=171 cells from 6 mice, P=0.0007) and KD MD mice (n=133 cells from 3 mice, p<0.0001) compared to WT mice (n=226 cells from 16 mice) (Kruskal-Wallis (KW= 45.64) followed by Dunn’s multiple comparisons post-hoc).

## Astroglial Cx30 closes the critical period by downregulating the RhoA-MMP9 pathway

We then sought to identify the molecular pathway through which Cx30 modulates PNN extent and maturation of PV interneurons. To this end we performed co-immunoprecipitation experiments with biotinylated WFA lectin followed by quantitative proteomics of pulled down proteins in WT and KD mice using label-free mass spectrometry. Reactome pathway analysis of enriched or unique proteins in WT compared to KD samples indicated an enrichment of proteins associated with the Rho GTPases-KTN1 pathway (Extended data Table 3, Fig. 4a upper panel). Conversely, analysis of proteins unique and enriched in KD compared to WT samples revealed enrichment of the RhoGTPases-ROCK pathway (Extended data Table 3, Fig. 4a). Besides, mass spectrometry analysis of Cx30 co-immunoprecipitates indicated interactions with Rho-GTPases signaling pathways, among which the Rho-associated protein kinase ROCK2 was significantly enriched in KD compared to WT (Extended data Table 4, Fig. 4a lower panel). Together, this exploratory approach points to a role of the Rho family of GTPases in the PNN changes between WT and KD mice.

**Figure 4.**
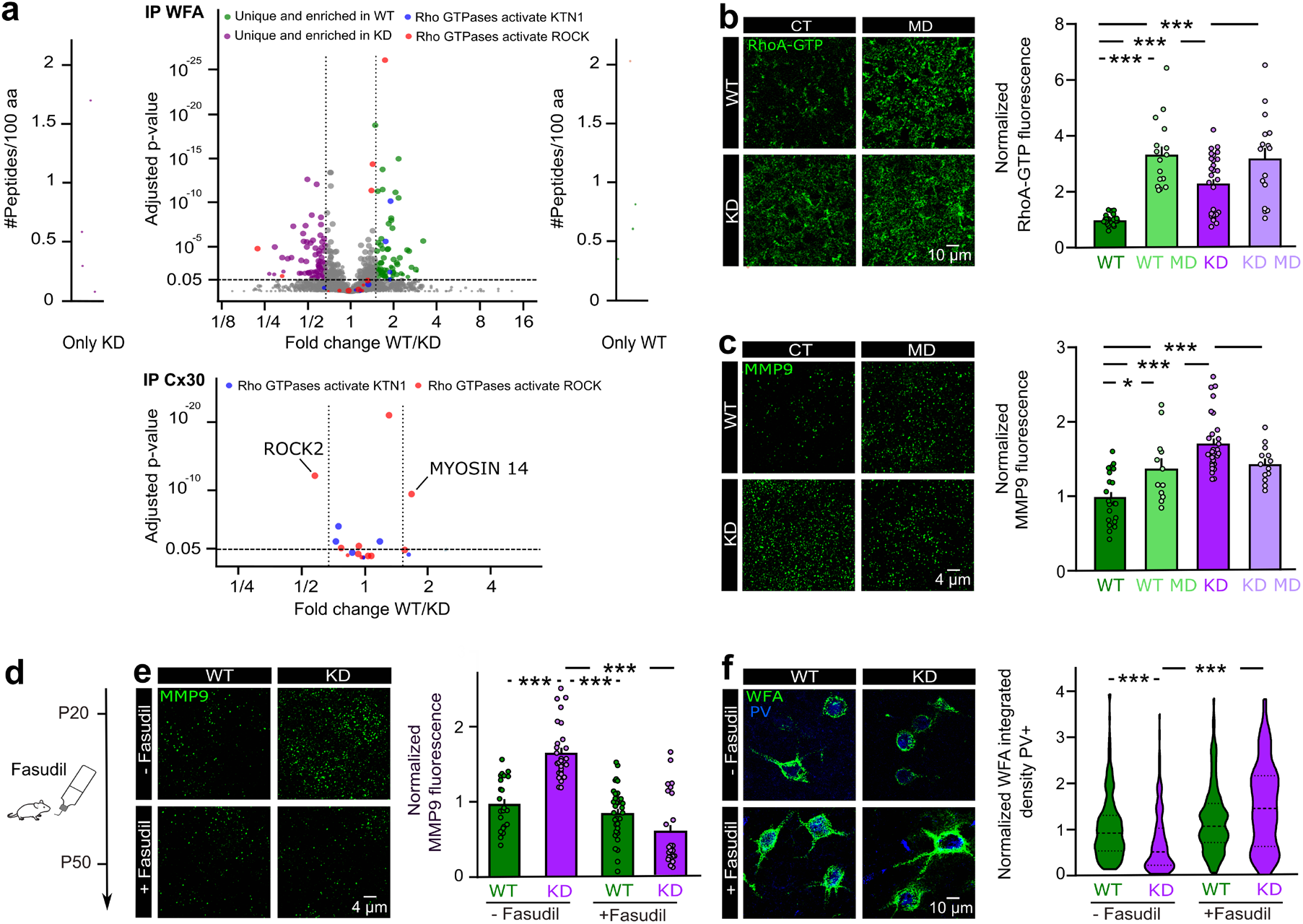
Astroglial Cx30 downregulates the RhoA-MMP9 pathway. (a) Top, volcano plot analysis representing the 2096 quantified proteins with at least 3 total peptides in all replicates enriched with biotinylated WFA lectin in KD and WT mice. Binding partners were obtained by using quantitative label-free mass spectrometry analysis performed from three replicates. Dashed vertical lines denote absolute fold change of 1.5 and the dashed horizontal line denotes the adjusted P-value of ratio significance of 0.05. Selected enriched proteins in WT (green) and KD (purple) samples are shown. Proteins from the enriched pathways Rho-GTPase activate KTN1 (blue) and Rho-GTPase activate ROCK (red) are highlighted. External plots show proteins with peptides identified only in one sample type (left in KD and right in WT). Bottom, volcano plot analysis representing interactors of Cx30 from the Rho-GTPase activate KTN1 (blue) and Rho-GTPase activate ROCK (red) pathways. The fold-change in WT (n=5) versus KD mice (n=4) are shown with selected proteins (Rock2 and Myosin 14) quantified with an absolute fold change ≥ 1.5, an adjusted P-value ≤ 0.05 and with ≥ 3 peptides. (b) Immunostaining for RhoA-GTP showed a marked increase in KD (n=27, P=0.0003), KD MD (n=16, P<0.0001) and WT MD mice (n=16, P<0.0001) compared to WT mice (n=22) (slices from 3 mice per group) (Kruskal-Wallis (KW=41.23) followed by Dunn’s post-hoc test. (c) Immunostaining for MMP9 shows a marked increase in KD mice (n=28, P<0.0001, DF=71), KD MD (n=13, P=0.0059, DF=71) and MD mice (n=12, P=0.0322, DF=71) compared to WT mice (n=22) (ANOVA (F(3,71)=15.07) followed by Tukey’s post-hoc test). (d) Diagram depicting the protocol for experimental treatment with Fasudil. (e) Fasudil rescued MMP9 levels in KD mice (KD+Fasudil: n=25 from 5 mice, KD=28 slices from 3 mice, P<0.0001), while it had no effect in WT mice (WT+Fasudil: n=34 from 6 mice, WT: 22 slices from 3 mice, P>0.9999, Kruskall-Wallis (KW=55.98) followed by Dunn’s post-hoc test). (f) Fasudil rescued perineuronal nets levels in KD mice (KD+Fasudil: n=52 from 5 mice; KD, n=177 from 16 mice, P<0.0001) while it had no effect in WT mice (WT+Fasudil, n=129 from 6 mice, WT, n=226 from 16 mice, P=0.1510) (Kruskall-Wallis (KW=71.65) followed by Dunn’s post-hoc test).

RhoA, one family member of the Rho-GTPases, plays a key role in cell remodeling^20^ through activation of ROCKs and its interactions with different connexins has been described in other cell types^21,22^. We thus tested the activation of RhoA by assessing levels of RhoA-GTP using immunohistochemistry, and found a strong increase in KD mice (Fig. 4b). In addition, while MD also increased RhoA-GTP in WT mice, it had no further effect in KD mice, suggesting that activation of RhoA pathway is a prerequisite for plasticity.

As the RhoA-ROCK pathway can modulate the expression of the ECM degrading enzyme Matrix Metalloproteinase-9 (MMP9)^23,24^, we next tested whether this signaling is involved in the Cx30 regulation of PNNs and PV interneurons maturation. To do so, we assessed MMP9 expression in layer 4 of the visual cortex, and found that MMP9 levels were markedly increased in KD mice compared to WT mice (Fig. 4c). In addition, akin to RhoA-GTP, while MD increased MMP9 levels in WT mice, it had no additional effect in KD mice (Fig. 4c). This suggests that activation of the RhoA-ROCK pathway leads to the degradation of PNNs through MMP9 in both KD mice and WT mice under MD condition. We next directly tested this hypothesis by inhibiting the ROCK signaling pathway in vivo with fasudil (Fig. 4d) and found that it rescued both MMP9 levels and PNN extent in KD mice, while it had no effect in WT mice (Fig. 4e, f). These data therefore indicate that astroglial Cx30 regulates MMP9 levels and PV cell maturation via the RhoA-ROCK signaling pathway.

## Discussion

We here demonstrate that astrocytes control timing for closure of the postnatal critical period of experience-dependent remodeling. We further identify that they achieve this through developmental rise of the astroglial protein Cx30, which we found to inhibit expression of MMP9 via the RhoA-ROCK pathway, thereby hindering maturation of local inhibition. While the latter is established as a trigger for critical period closure, its control by astroglial signaling place astrocytes as key elements orchestrating the shaping of synaptic circuits. Beyond its role in critical period, the identified pathway may well be involved in structural synaptic plasticity associated with cognitive functions, since we previously found that Cx30 regulates the structure and efficacy of the tripartite synapse in the hippocampus^25^. The regulation of MMP9 levels that we here describe occurs via an unconventional signaling through Cx30 and thus represents a novel astroglial pathway regulating wiring of brain circuits. Because extended critical periods are associated with neurodevelopmental defects resulting in sensori-motor or psychiatric disorders^4,5^, these findings provide a new target for the development of strategies aiming at re-inducing a period of enhanced plasticity in adults and favor rehabilitation after brain damage or developmental malfunction.

## Acknowledgments

This work was supported by grants from the European Research Council (Consolidator grant #683154), European Union’s Horizon 2020 research and innovation program (Marie Sklodowska-Curie Innovative Training Networks, grant #722053, EU-GliaPhD) and UNADEV-Aviesan to N.R., from Région Ile-de-France and Fondation pour la Recherche Médicale to D.L., from College de France to G.D., from Fyssen Fondation to J.Z. and from French Research Ministry (Biosigne Doctoral school) to R.B. The authors thank Noëlle Dufour, Charlène Joséphine and MIRCen's viral vector facility, the animal and imaging facilities of College de France as well as Michel Roux from the mouse clinical institute for excellent technical assistance.

## Author Contributions

Conceptualization: N.R., G.D., J.R. C.M.; Data curation: G.D., J.R., R.B. D.L., N.R.; Formal analysis: G.D., J.R., R.B., P.B., J.Z., V.S., D,L., N.R.; Funding acquisition: N.R., G.D.; Investigation: J.R., R.B., G.D., C.F.C., F.D., P.E., K.S., J. Z., N.R.; Methodology: J.R., G.D., J.M.; Resources: D.L., A.P.B.; Supervision: N.R., G.D.; Project administration: N.R.; Validation: J.R., G.D., N.R.; Visualization: J.R., R.B., G.D., N.R.; Writing original draft: G.D., N.R.; Writing – review & editing: all authors.

## Additional Information

**Supplementary Information** is available for this paper.

**Correspondence and requests for materials** should be addressed to N.R.

## Extended Data

**Extented Table 1.**
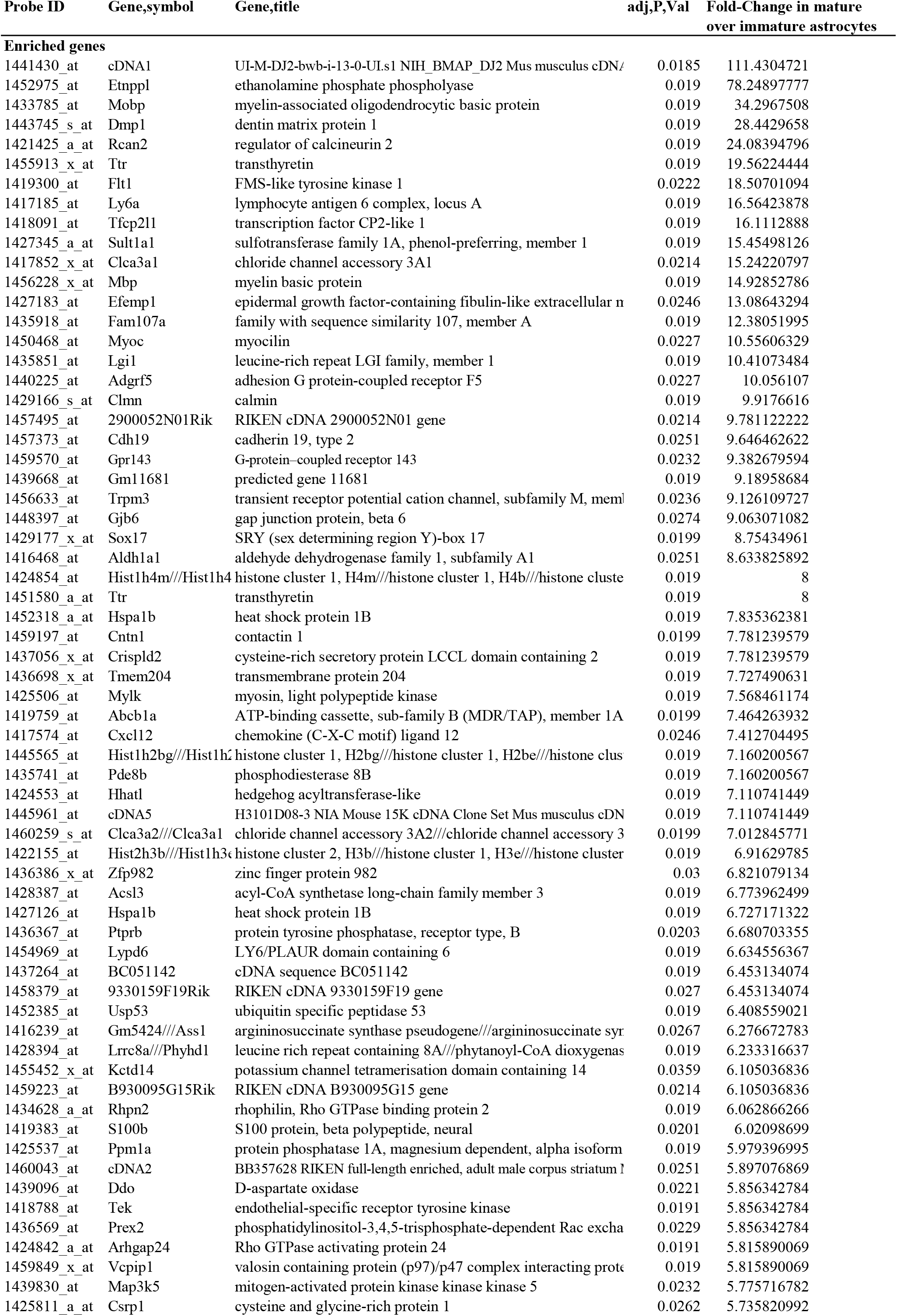
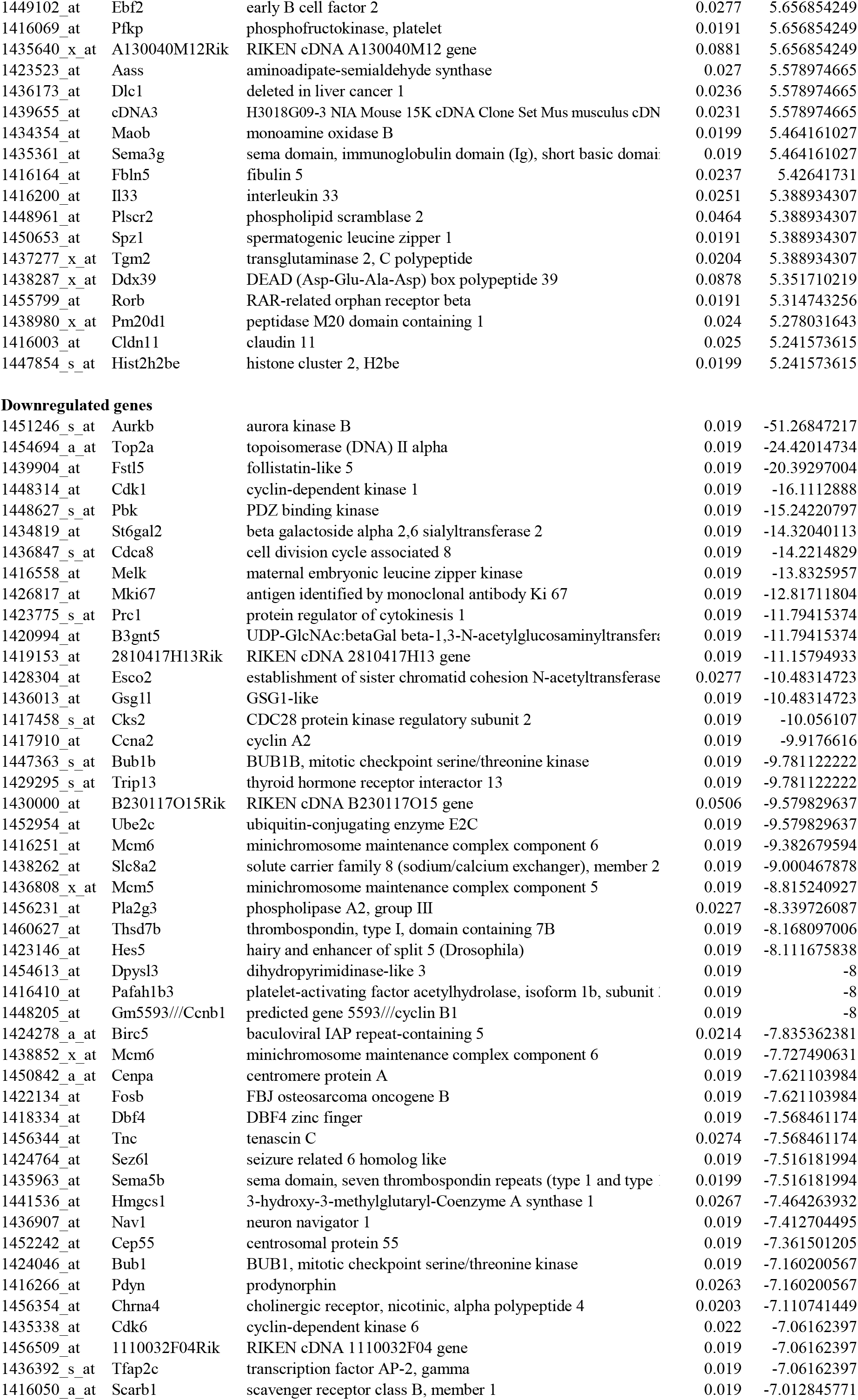
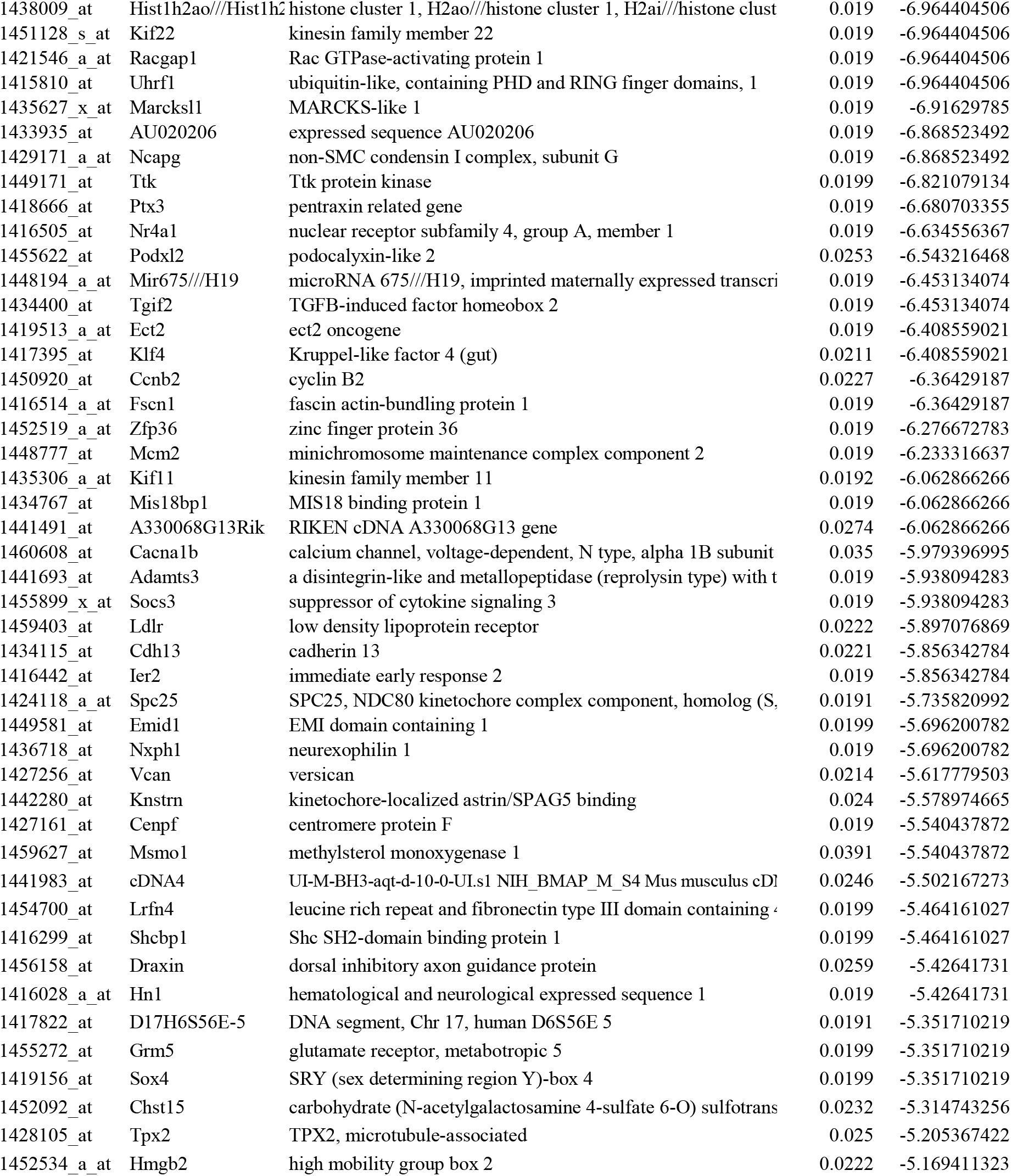
Genes statistically enriched and down-regulated in developing astrocytes. The differentially expressed genes in mature astrocytes (P30) compared to immature astrocytes (P7) are listed along with their fold-change enrichment and the Affymetrix probe set ID (Cahoy et al., 2008). All genes are statistically different by Geo2R analysis with an adjusted P-value (Benjamini and Hochberg, 1995 FDR) threshold <5%. There are 93 and 82 immature and mature astrocyte-enriched genes, respectively, selected with a 5-fold change criteria.

**Extented Table 2.**
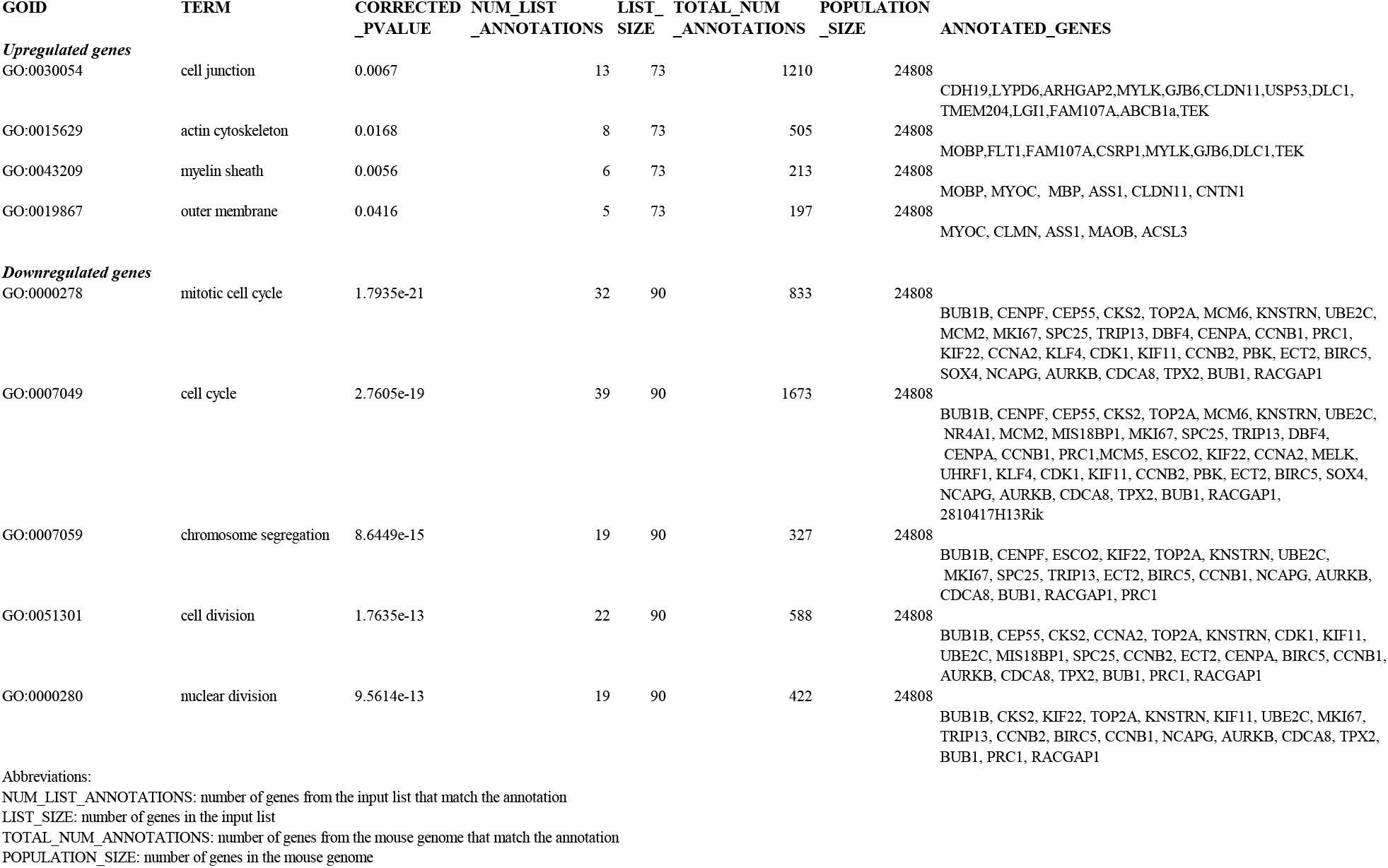
GO Term Finder analysis of genes upregulated and downregulated in mature vs immature astrocytes. Selected lists of differentially expressed genes (93 down-and 82 up-regulated genes) were analyzed for GO terms selecting “process” for immature associated genes and “component” for mature ones. Whereas immature astrocytes are characterized by intensive cell division, mature astrocytes present enrichment in genes involved in cell junction.

---

**Extented Figure 1.**
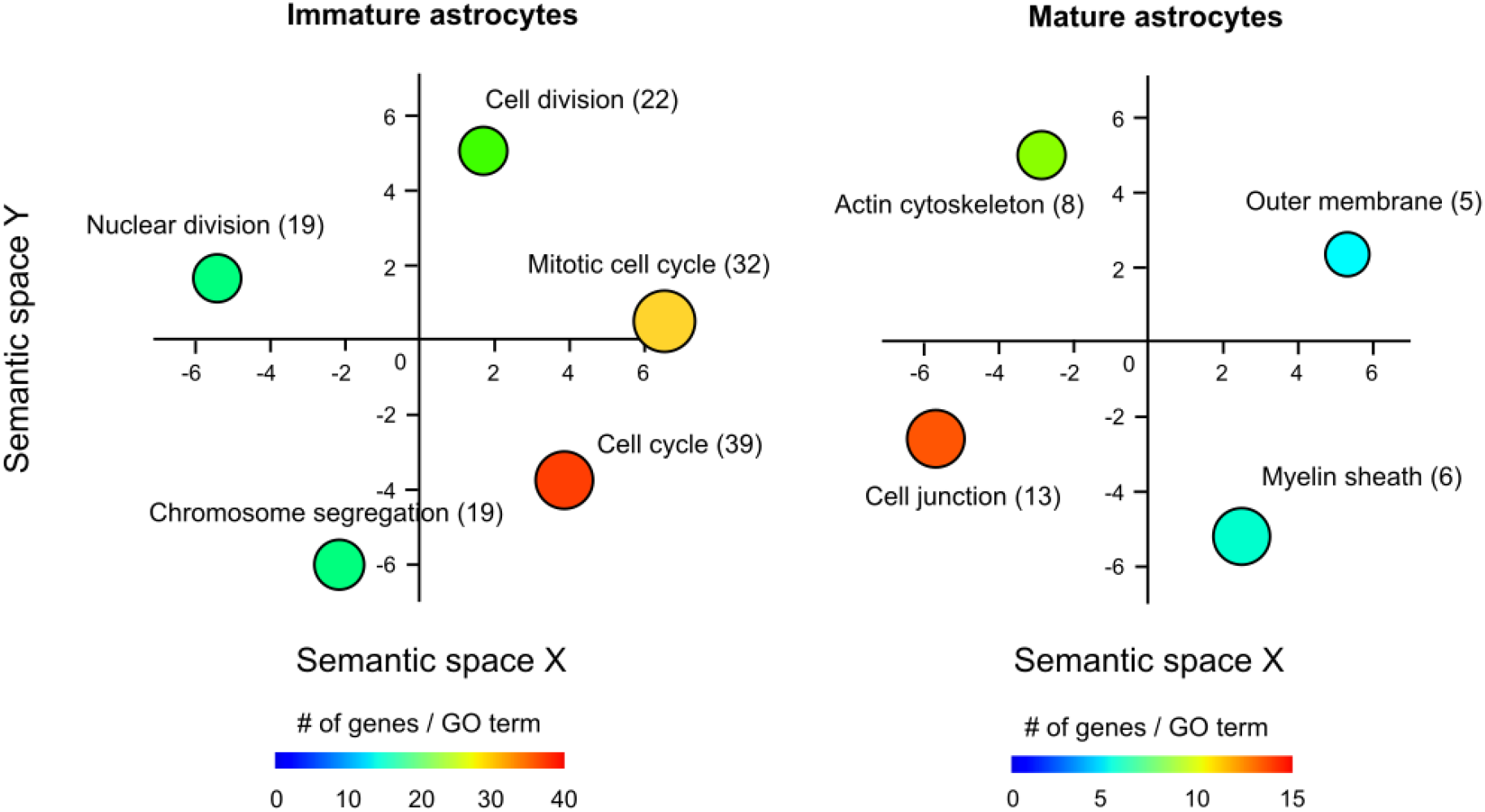
Semantic similarity-based scatterplots of GO terms. GO terms (Extended Data Table 2) identified through the analysis of down or up-regulated genes (Extended data Table 1) respectively associated to immature (P7) and mature (P30) astrocytes and visualized according to the similarity of the GO terms along arbitrary X and Y axis. Each circle is color-coded according to the frequency of the GO term in the EBI GOA reference database, and the number of genes per GO term is indicated in parenthesis. Circle size indicates the P-value (circles with bigger size have smaller P value) (Extended Table 2).

**Extented Figure 2.**
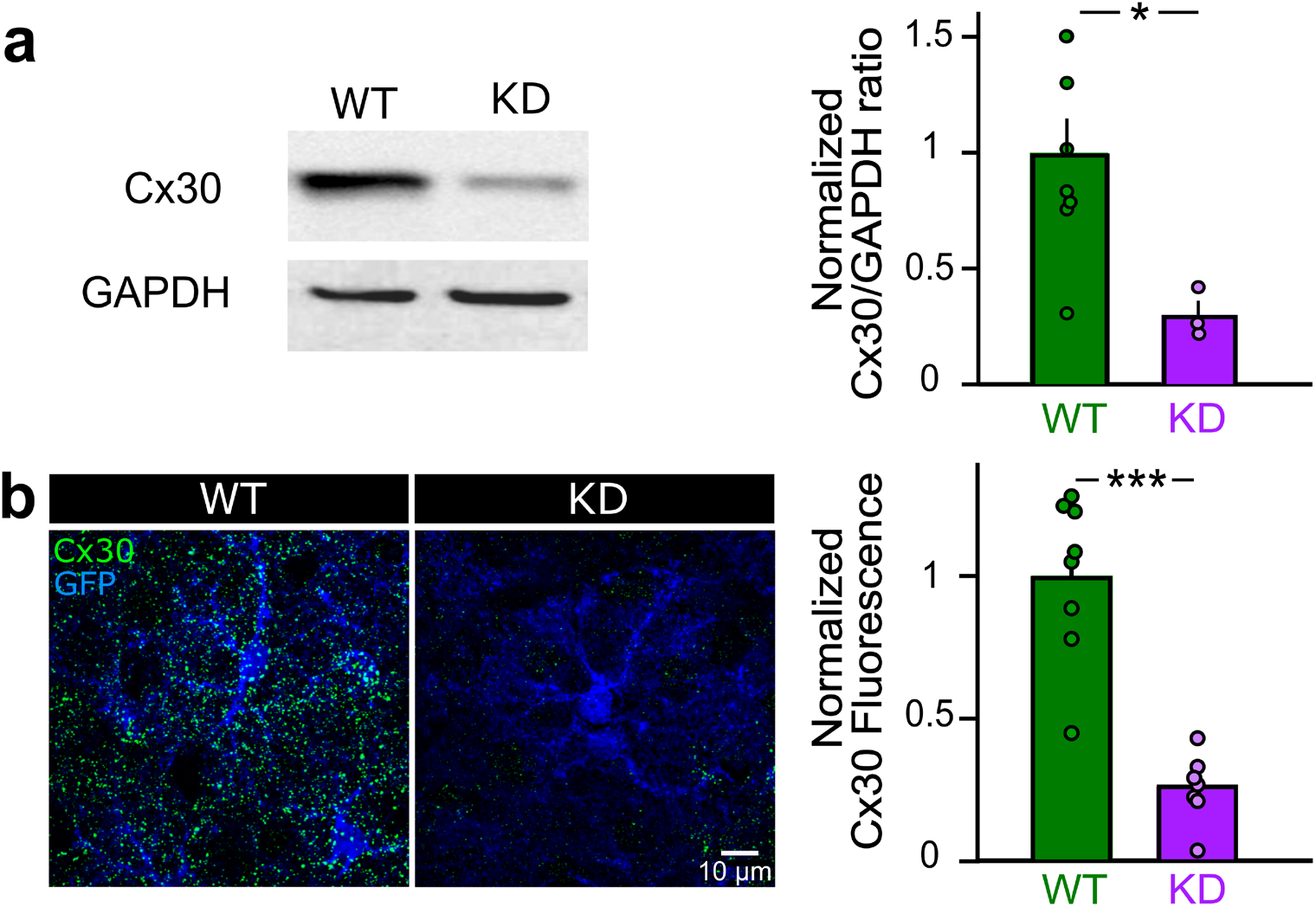
Characterization of Cx30 KD mice. (a) Western blot analysis from visual cortex shows a decrease of 70% in Cx30 expression in KD mice (n=3) compared to WT mice (n=8, P=0.0242, U=1, Mann-Whitney test). (b) Immunostaining of Cx30 in the visual cortex also shows a 70% decrease in Cx30 expression in astrocytes in KD mice (n=7 samples from 3 mice) compared to WT mice (n=8 samples from 3 mice, P=0.0003, U=0, Mann-Whitney test).

**Extented Figure 3.**
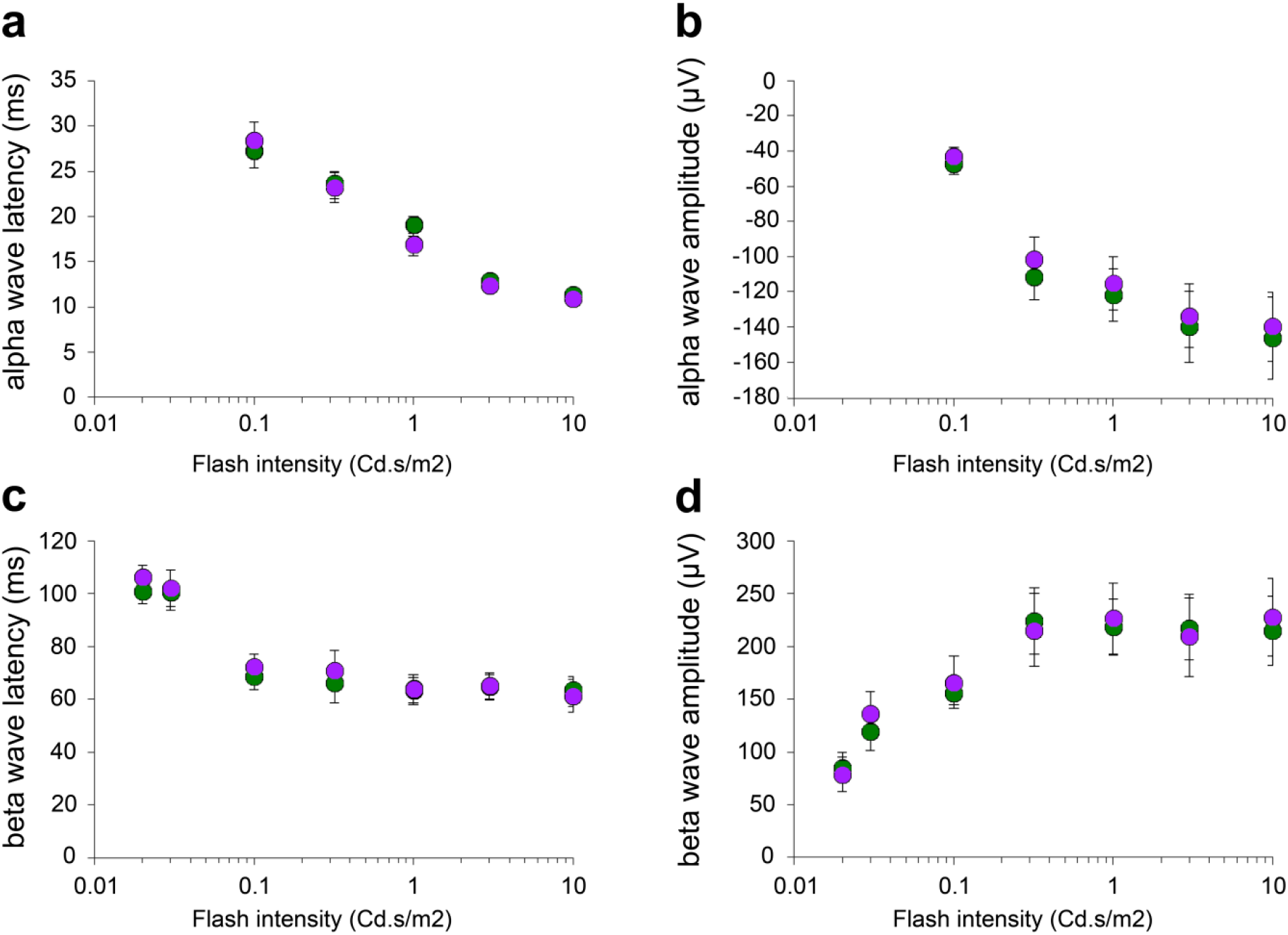
Retinal activation is unchanged in KD mice. The changes in potentials induced by light flashes of increasing intensity (from 0.02 to 10 Cd.s/m2) in the retinal tissue was assessed by recording alpha- and beta-wave latency and amplitude. No difference in electroretinograms were detected between WT (n=6) and KD mice (n=6, alpha wave latency: P=0.5550, DF=50, F(1,50)=0.3531; alpha wave amplitude: P=0.6231, DF=50, F(1,50)=0.2446; beta wave latency: P=0.5278, DF=71, F(1,71)=0.4027; beta wave amplitude: P=0.8016, DF=70, F(1,70)=0.06365; 2-way ANOVA).

**Extented Table 3.**
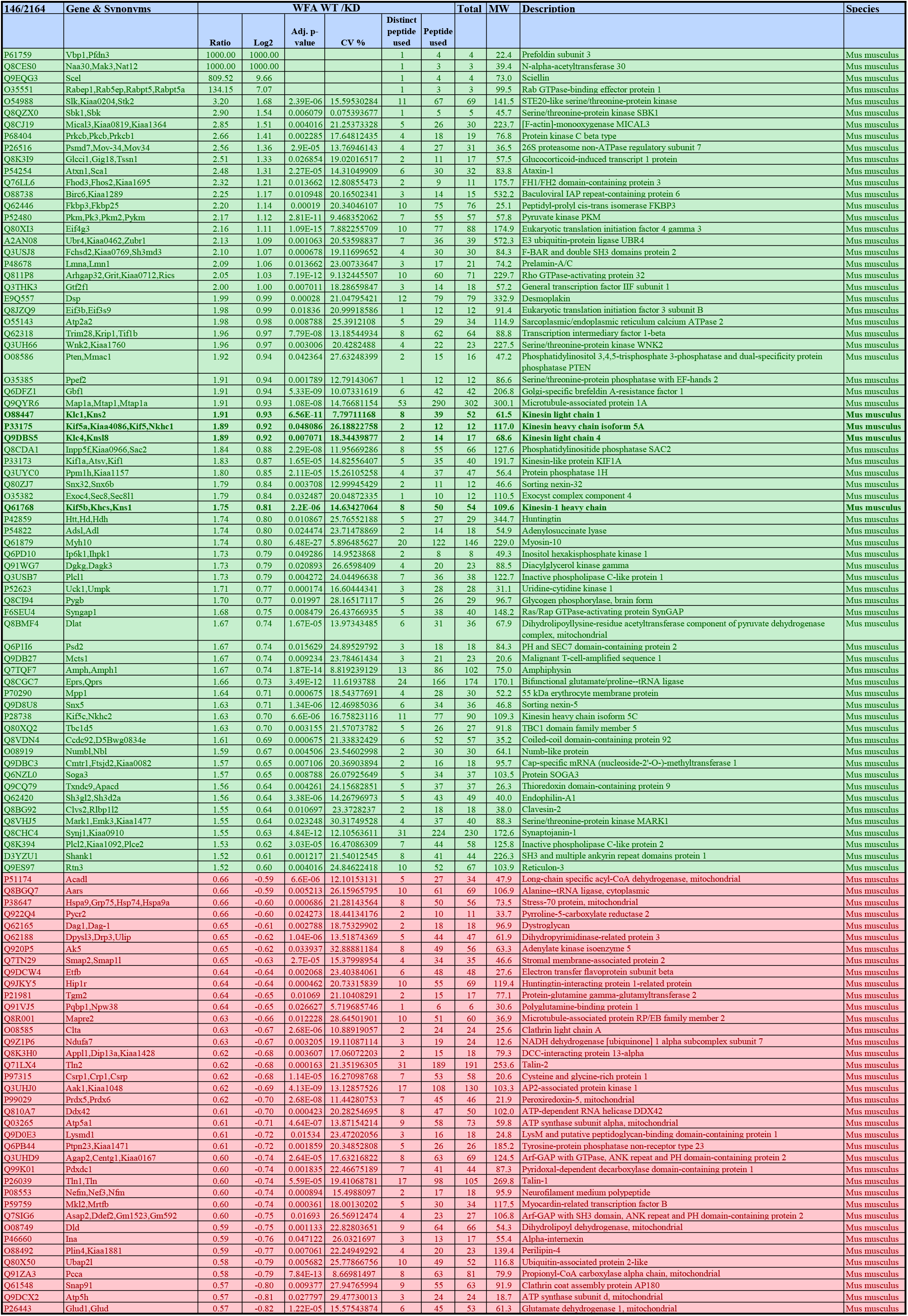
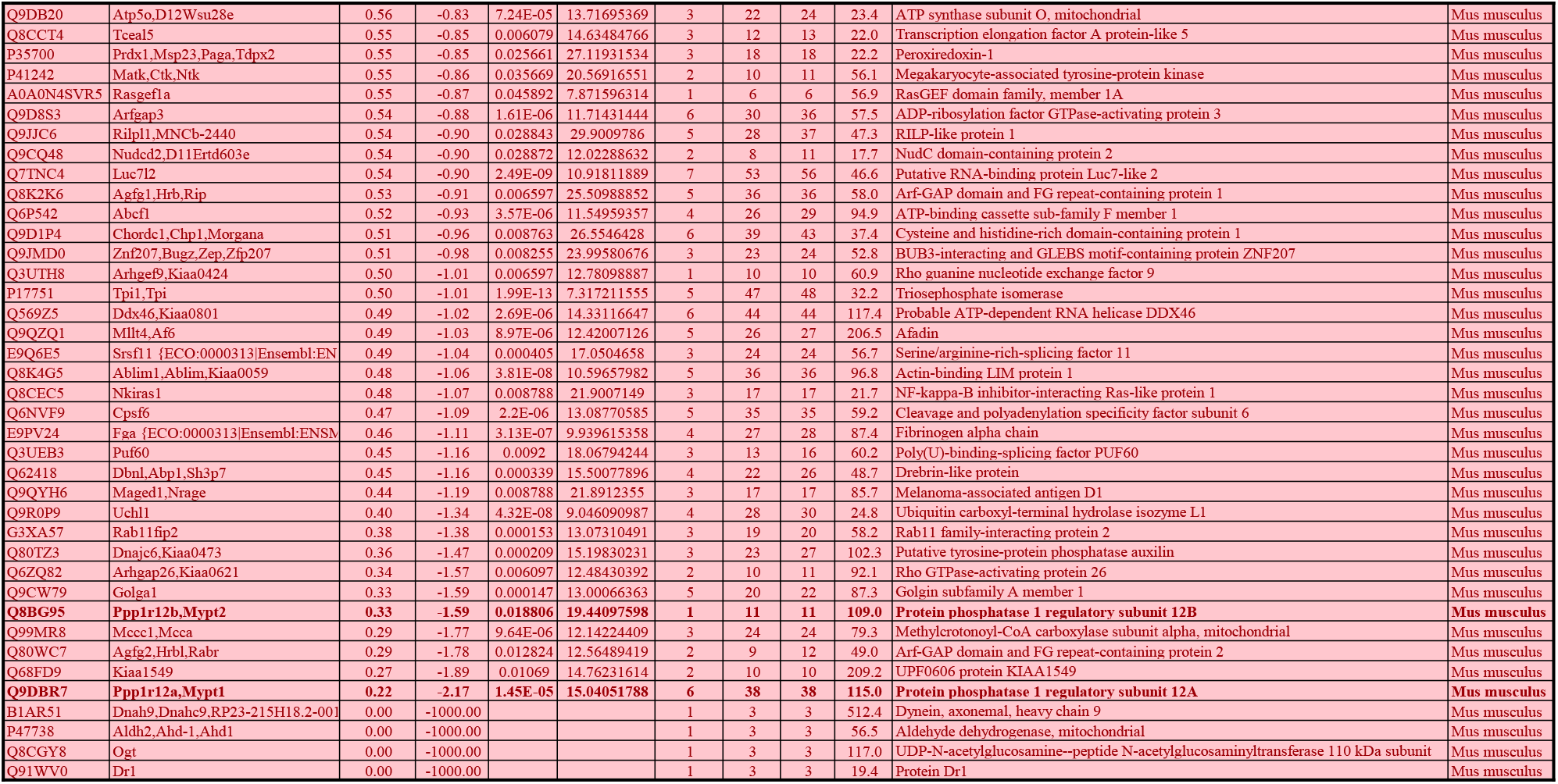
List of the WFA protein partners. Analysis of the changing WFA interactors in WT vs KD mice. In WT cells, 70 proteins were enriched or unique (log2 ratio=1000) compared to Cx30 KD cells (with the following parameters: number of peptides≥3, ratio≥1.5 and p-value≤0.05), while in Cx30 KD cells 77 proteins were enriched or unique (log2 ratio = −1000) compared to WT cells (with the following parameters: number of peptides≥3, ratio≥1/1.5 and p-value≤0.05). WT and KD proteins are highlighted in red and green, respectively. Protein from the pathways Rho GTPase activate KTN1 (bold green) and Rho GTPase activate ROCK (bold red) are indicated.

**Extented Table 4.**
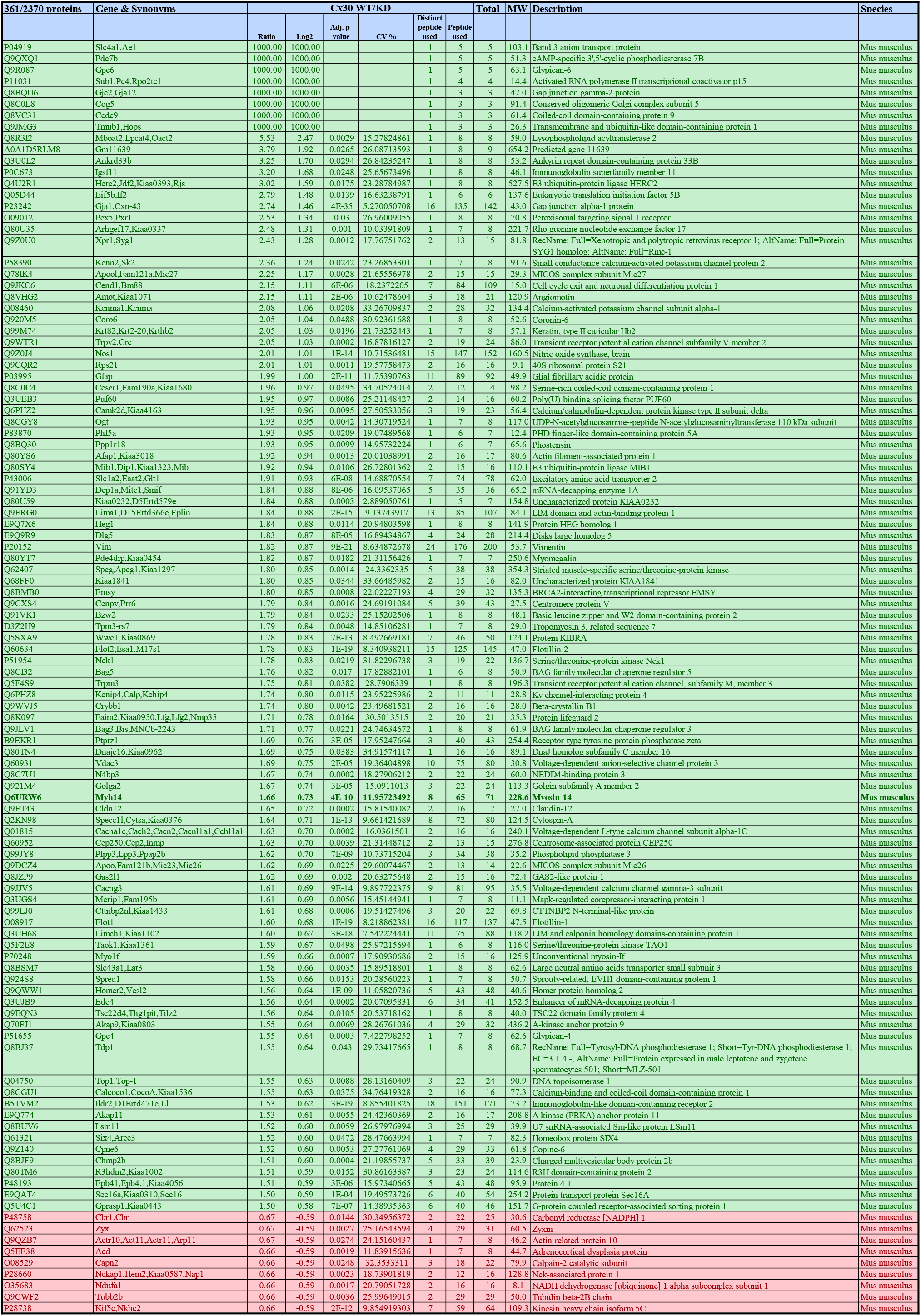
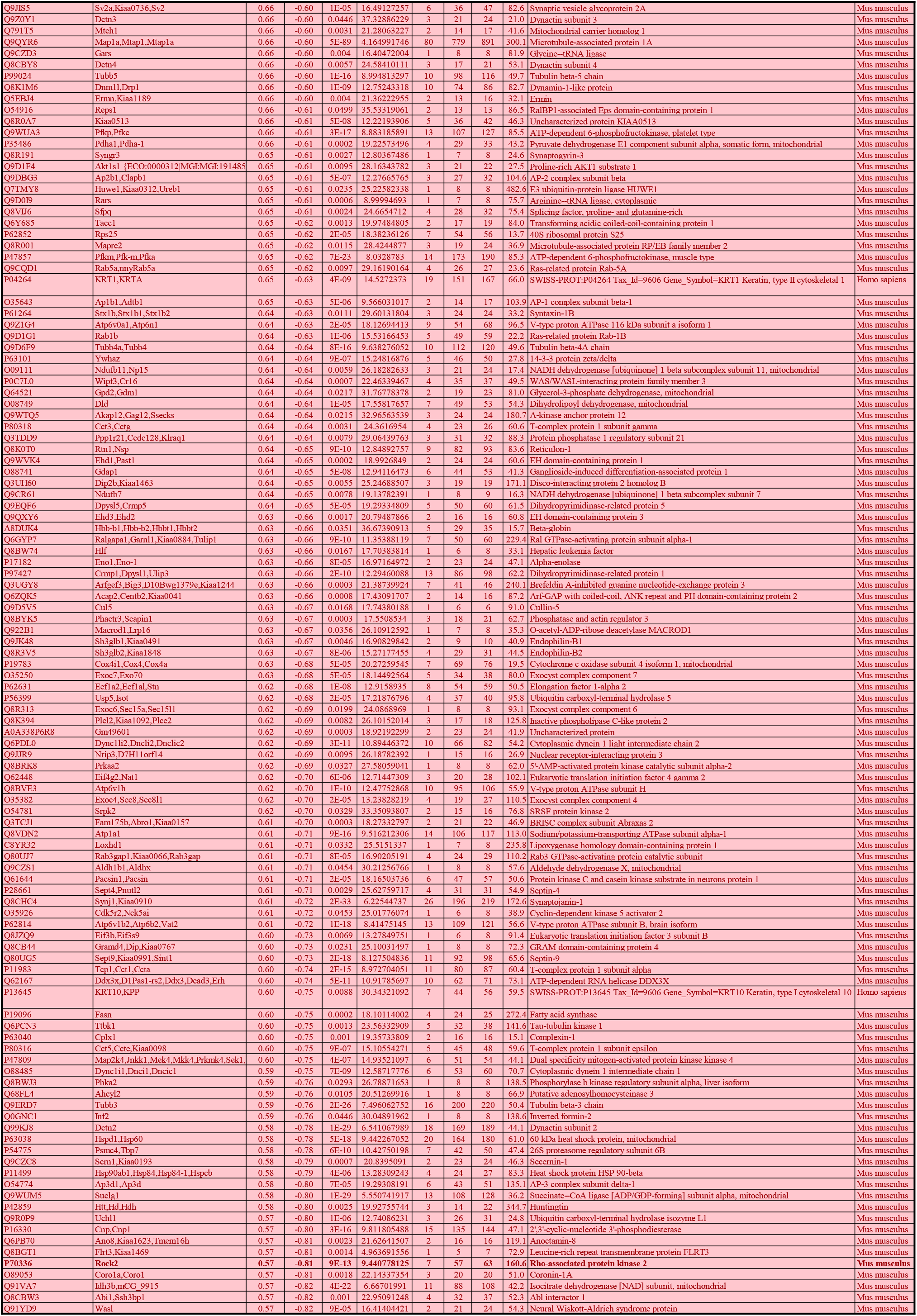
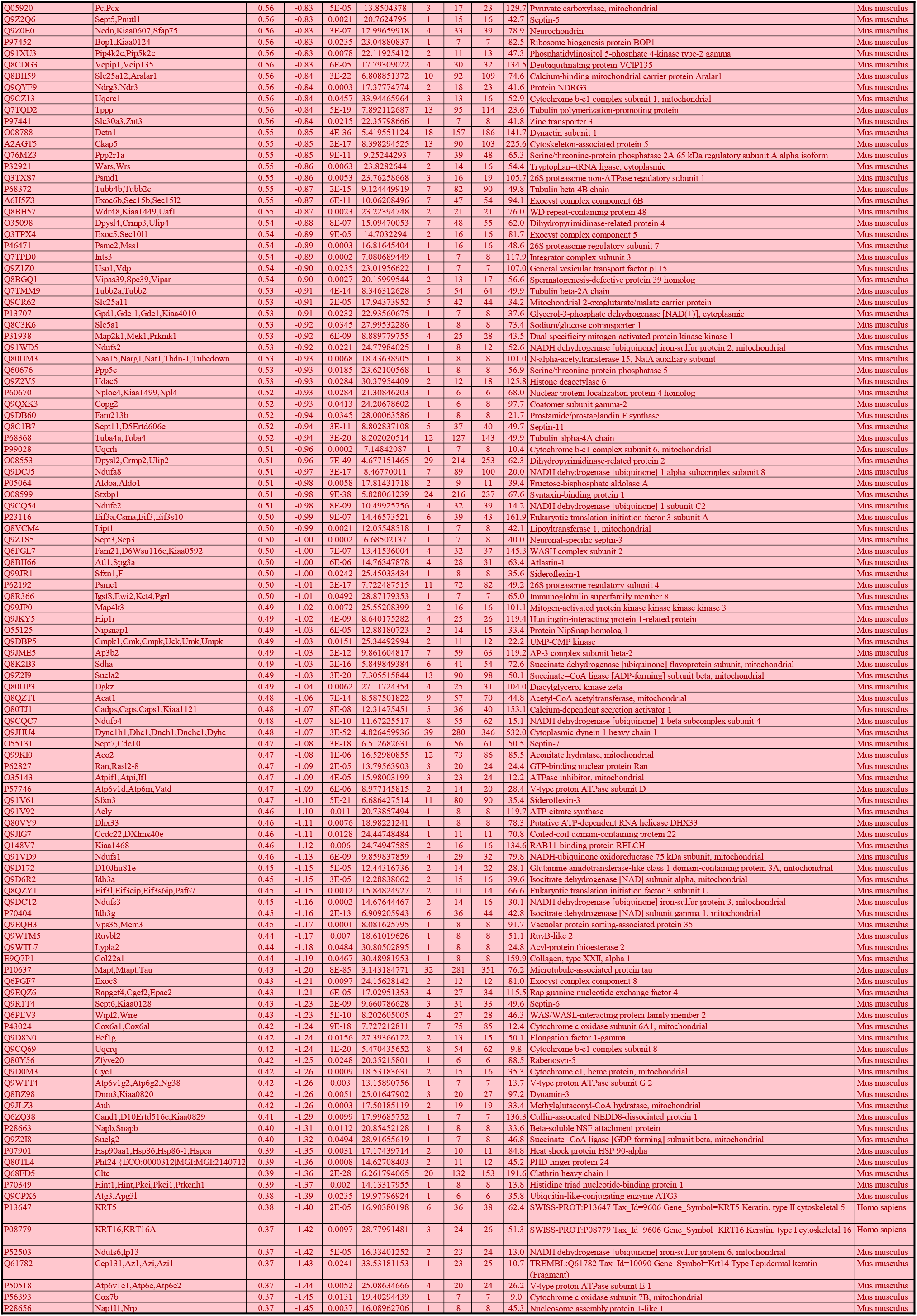
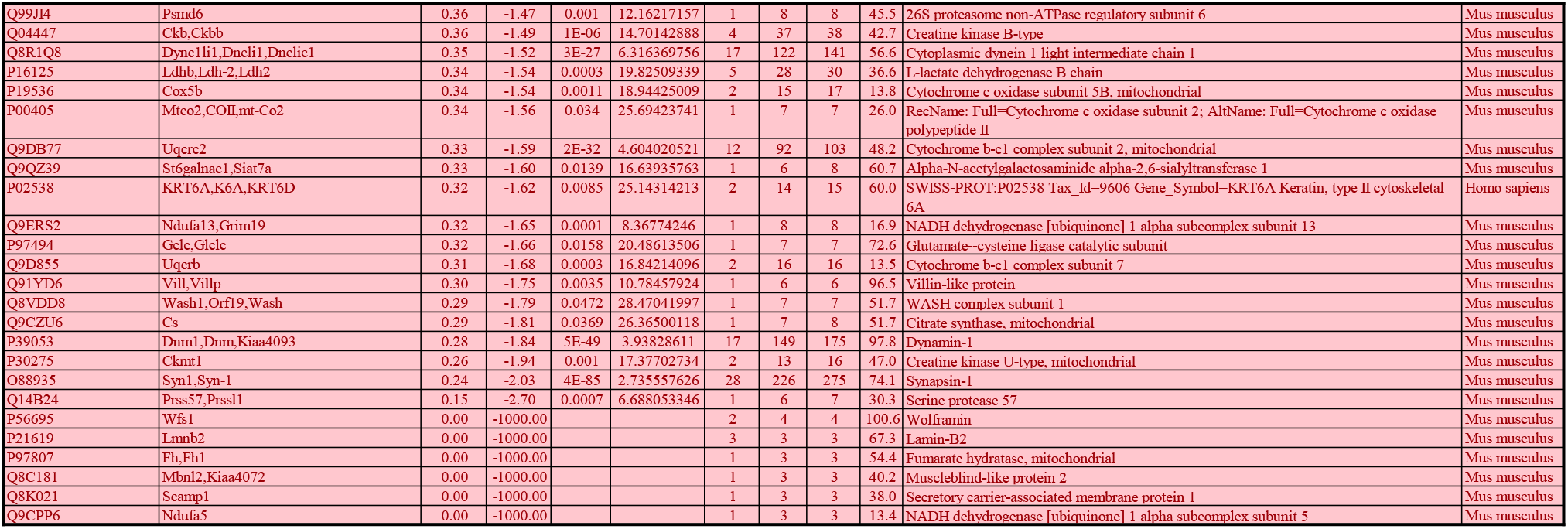
List of the Cx30 protein partners. Analysis of the changing Cx30 interactors in WT vs KD mice. In WT cells 100 proteins were enriched or unique compared to KD cells (with the following parameters: number of peptides ≥ 3, ratio ≥ 1.5 and p-value ≤ 0.05), while in Cx30 KD cells, 261 proteins were enriched or unique (log2 ratio = −1000) compared to WT cells (with the following parameters: number of peptides ≥ 3, ratio ≥ 1/1.5 and P-value ≤ 0.05). WT and KD proteins are highlighted in red and green, respectively. Proteins from the pathway Rho-GTPase activate ROCK are indicated for WT (bold green, Myosin 14) and KD (bold red, Rock2) samples.

## Supplementary Information

### Methods

#### Animals

All procedures on animals were performed according to the guidelines of European Community Council Directives of 01/01/2013 (2010/63/EU) and our local animal welfare committee (Center for Interdisciplinary Research in Biology in College de France, certificate A751901, Ministère de l’Agriculture et de la Pêche). Experiments were carried out using mice of wild type (WT) C57BL/6j background, mice expressing enhanced green fluorescent protein under the astrocytic promoter aldehyde dehydrogenase 1 family member L1 (Aldh1l1-eGFP)^1^ (JAX stock #026033), as well as constitutive knockout mice for Cx30 which were previously characterized^2^ and an astroglial conditional knockdown mouse line for Cx30 that we generated (hGFAP-Cre-Cx30^fl/fl^, named KD). Animals were group housed on a 12 h light/dark cycle. All mice were backcrossed to the C57BL/6J background. Mice of both genders and littermates were used. All efforts were made to minimize the number of animals used and their suffering.

#### Generation of Cx30 knockdown mice

Astroglial Cx30 knockdown mice (KD) were generated by crossing the hGFAP-cre line constitutively expressing the cre recombinase transgene driven by the human astrocytic glial fibrillary acidic protein (hGFAP) promoter^3^ (JAX stock #004600) with Cx30^fl/fl^ line containing cre-excisable loxP sequences in the endogenous *Gjb6* gene^2^.

#### Primary astrocytes cultures

Primary cortical astrocyte cultures were prepared as previously described^4^. Briefly, brains were removed (either P1-P3 pups, or P19 mice) and caudal cortices (therefore enriched in visual cortex material) were dissected in cold PBS-glucose (33 mM). Meninges were carefully removed and P1-P3 cortices were mechanically dissociated with Pasteur pipette in PBS-glucose to obtain single cell suspension. For P19 animals, caudal cortices were dissected, treated with papain (0.8 mg/ml; Worthington) for 30 min at 37°C and washed, then mechanically dissociated in PBS containing 5% FCS, 0.3% glucose and 5mM HEPES to obtain a single cell suspension. These dissociated single cell preparations were transduced with a HIV-1-derived lentivirus vector expressing GFP under the control of the PGK ubiquitous promoter (LV-PGK-GFP). Lentiviral particles were produced by transient transfection of HEK-293T cells with a three plasmids system and vector titers were quantified using an ELISA assay of p24 antigen (Gentaur, France), both as previously described^5^. Infection was performed in DMEM containing 5% SVF, 100ng of viral particles were added to 10^6^ cells for 3 hours at 37°C. The suspension of infected cells was then directly seeded on polyornithine-coated glass coverslips or petri dishes (0.1 mg/ml) in DMEM containing 10% heat-inactivated FCS, 100 U/ml penicillin/streptomycin (GIBCO) and incubated at 37°C, 5% CO2. 24 hours after transduction, the medium was replaced. After one week, once cells have reached confluency, 1 μM of cytosine-beta-D-arabinofuranoside was added to the cell culture for 2 days, to eliminate proliferating microglial cells. Medium was then changed every 3 days and astrocytes were used after 10 days in culture (10DIV). For immunocytochemistry, cells were fixed with 4% PFA and washed twice in PBS before proceeding with immunostaining. For intracerebral injection, the culture was washed with PBS, then incubated with trypsin 0.25 EDTA% (Invitrogen) for 5 to 10 min at 37°C. The harvested cells were collected in fresh DMEM 5% SVF and incubated for 2 hours at 37°C, to allow cell recovery.

#### Recombinant adeno-associated virus (rAAV) generation

For rAAV *in vivo* gene transfer, a transgene composed of GFP cDNA was placed under the control of a GFAP-specific promoter in a rAAV shuttle plasmid containing the inverted terminal repeats (ITR) of AAV2 (AAV-GFAP-GFP). Pseudotyped serotype 9 rAAV particles were produced by transient co-transfection of HEK-293T cells, as previously described^6^. Viral titers were determined by quantitative PCR amplification of the ITR on DNase-resistant particles and expressed as vector genome per ml (vg/ml).

#### >Stereotaxic intracerebral injections

Animals were anaesthetized i.p. with a mix of ketamine (95mg/kg) and xylazine (12mg/kg) in 0.9 % NaCl and fitted into a stereotaxic frame (David Kopf Instruments). For cells injection, a 2μl-gauge Hamilton syringe adapted to KDS100 pump (Kd Scientific) was linked to a glass capillary (Harvard apparatus, 30-0041) that was surgically implanted in the right visual cortex area with the following coordinates to the Lambda: antero-posterior 0, medio-lateral + 2.8 and dorso-ventral - 0.5 mm. A maximum of 1.5μl of cell suspension (20×10^4^ cells) in DMEM containing 0.25% SVF was infused at a flow rate of 0.2 μl/min. Control animals were injected the same way with 1.5μl of medium. After the injection, the capillary was left in place for 5 min and then slowly withdrawn.

Alternatively for *in vivo* cell labeling, 1μl of rAAV2/9-GFAP-eGFP (diluted in PBS at a concentration of 1×10^10^ vg/μl) was unilaterally injected in the right visual cortex at the rate of 0.2μl/min using the same coordinates from Bregma mentioned above. The injection was performed using 29-gauge blunt-tip needle connected to 2μl Hamilton syringe and the injection rate was controlled by syringe pump (KD Scientific). After the injection, the needle was left in place for 5 min and then slowly withdrawn. Following surgery, mice were allowed to recover from anesthesia on a heating pad and monitored for the next 24h.

#### Gene Ontogy and enrichment analyzes of transcriptomic data

Comparing microarray data from P30 to P7 astrocytes (GEO#: GSE9566/plateform#: GPL1261) using Geo2R analyzes (https://www.ncbi.nlm.nih.gov/geo/info/geo2r.html)^7^ allowed to identify differentially expressed genes between the two stages. Using a cut-off of 5 fold-change, about 200 up and down-regulated genes were analyzed for enrichment using their Gene Ontology assignment (GOtermFinder: https://go.princeton.edu/cgi-bin/GOTermFinder?). Process and Component GO options were selected respectively for the down and up DEG groups. Finally GO results were Visualized using ReViGO (http://revigo.irb.hr/).

#### Antibodies, immunohistochemistry and immunoblotting

All the antibodies used in this study are commercially available and have been validated in previous studies, as reported by the suppliers. The following primary antibodies were used: Cx30 rabbit polyclonal (1:500, 71-2200, Zymed), Cx43 mouse monoclonal (1:500, 610061 BD Biosciences), GFAP mouse monoclonal (1:500, G3893, Sigma-Aldrich), chick anti GFP (1:500, AB13970, Abcam), Parvalbumin mouse monoclonal (1:500, 235, SWANT), MMP9 rabbit (1:500, 3852, cell signaling), anti-active RhoA mouse monoclonal (1:500, 26904, NewEast Biosciences), Lectin from Wisteria Floribunda, biotin conjugate (1:500, L1516-2MG, Sigma-Aldrich). The following fluorescent dye-conjugated secondary antibodies were used in appropriate combinations: goat anti-mouse IgG conjugated to Alexa 555 (1:2000, A-21424, Thermo Fisher), goat anti-rabbit IgG conjugated to Alexa 647 (1:2000, A-21245 Thermo Fisher). goat anti-chicken Ig conjugated to Alexa 488 (1:2000, A-11039, Thermo Fisher), Streptavidin Alexa fluor 488 (1:2000, S11223, Molecular Probes).

Immunohistochemistry and quantifications were performed as follows. Briefly, animals anesthetized with lethal dose of Dolethal (150μl/10g), were perfused by intracardiac with PBS first and 2% paraformaldehyde (PFA). The brains were carefully removed for an overnight post-fixation in the same fixative, followed by transfer in 30% sucrose for cryoprotection. Brain coronal microtome sections (40μm thick) were collected in PBS and pre-incubated 1 h with PBS-1% gelatin in the presence of 0.25% Triton-X100 (PGT). Brain sections were then stained overnight at 4°C with primary antibodies and washed in PGT three times. Appropriate secondary antibodies with DAPI (1:200, D9564, Sigma-Aldrich) were finally applied for 2 hours at room temperature. After several washes in PBS, brain slices were mounted in Fluoromount (Clinisciences) and examined with an inverted confocal laser-scanning microscope (Confocal Leica SP5 inverted). Stacks of consecutive confocal images taken with a 63x objective at 600-1000 nm intervals were acquired sequentially with two lasers (argon 488 nm and helium/neon 543 nm or 647 nm) and Z projections were reconstructed using image J software. Alternatively, primary astrocyte cultures were fixed at room temperature with 2% paraformaldehyde for 10 min, washed twice with PBS and incubated 1h with 5% non-immune goat serum (Zymed) in the presence of 0.25% Triton-X100 before proceeding as described above.

Quantifications of RhoA-GTP, WFA, MMP9, and Cx30 expression were performed with ImageJ software. WFA integrated density was determined using histogram-based thresholding method specifically around PV interneurons. RhoA-GTP and MMP9 fluorescence were evaluated by measuring the global mean intensity or the integrated density, respectively. The integrated density of Cx30 was quantified specifically in astrocytes labeled with GFP following infection with rAAV2/9-GFAP-eGFP, as described above. For the quantification of WFA, Aldh1l1 and Cx30 staining across cortical layers, confocal image stacks were acquired on four to five brain sections from each animal. Regions of interest (200 x 630μm) were randomly defined in V1 and gray values were averaged along the transversal axis (bin 1.5μm). Western blotting and quantification were performed as previously described^8^. Shortly, cells were collected with a folded pipette tip (200μl) in a small volume of PBS containing protease inhibitor cocktail (Euromedex), phosphatase inhibitors (Beta-glycerophosphate, 10 mM) and orthovanadate (1 mM), to which Laemmli 5X buffer was added. Alternatively, visual cortex samples were isolated from acute slices of 400μm cut in cold ACSF 1X. The same protocol was applied for visual cortex tissue using SDS 2% instead of PBS. Samples were sonicated, boiled 5 min and loaded on 4-12% polyacrylamide gels. Proteins were separated by electrophoresis and transferred onto nitrocellulose membranes. Membranes were saturated with 5% fat-free dried milk in triphosphate buffer solution and incubated overnight at 4°C with primary antibodies. They were then washed and exposed to HRP-conjugated secondary antibodies: goat anti rabbit IgG (1:2500, CSA2115 Clinisciences), goat anti-mouse IgG (1:2500, CSA2108). The HRP-conjugated primary anti-GAPDH antibody (1:10,000, G9295 Sigma-Aldrich) was used as loading control. Specific signals were revealed with the chemiluminescence detection kit (Western Lightning plus-ECL, NEL103E001EA, Perkin Elmer). Semi-quantitative densitometric analysis was performed after scanning the bands with the image J software.

#### Proteomics and Mass Spectrometry Analysis

##### Sample Preparation

Visual cortices of WT and Cx30KD mice were dissected from acute slices of 400 μm cut in cold ACSF 1X with a vibratome (Leica VT1200S). Samples were prepared in a lysis buffer containing 0.32M sucrose, 5 mM HEPES, 10 mM MgCl_2_ and complete EDTA free, and centrifuged (8 min at 1700 g) at 4°C. The supernatant was incubated overnight with lectin from Wisteria Floribunda Agglutinin, biotin conjugate (1:500, L1516-2MG, Sigma-Aldrich) and anti-Cx30 biotinylated (using anti-Cx30 rabbit polyclonal 71-2200, Zymed and Biotinylation kit/Biotin conjugation kit ab201795, abcam) or - coupled with streptavidin beads (Immunoprecipitation Kit - Dynabeads Protein G, 10007D, Invitrogen). Then, the beads were washed with lysis buffer before mass spectrometry analysis. Two additional washes in 100 μL of ABC buffer (25 mM NH4HCO3) were performed keeping the beads in the magnet and with no incubation. Finally, beads were resuspended in 100 μl of ABC buffer and digested by adding 0.20 μg of trypsine/LysC (Promega) for 1 hour at 37 °C. Samples were then loaded onto homemade Tips packed with Empore™ C18 Extraction Disks (3M™ Discs 2215) for desalting. Peptides were eluted using 40/60 MeCN/H2O + 0.1% formic acid and vacuum concentrated to dryness.

##### LC-MS/MS Analysis

For the biotinylated WFA lectine pull down, liquid chromatography (LC) was performed with an RSLCnano system (Ultimate 3000, Thermo Scientific) coupled online to a Q Exactive HF-X with a Nanospay Flex ion source (Thermo Scientific). Peptides were first trapped on a C18 column (75 μm inner diameter × 2 cm; nanoViper Acclaim PepMapTM 100, Thermo Scientific) with buffer A (2/98 MeCN/H2O in 0.1% formic acid) at a flow rate of 2.5 μL/min over 4 min. Separation was then performed on a 50 cm x 75 μm C18 column (nanoViper Acclaim PepMapTM RSLC, 2 μm, 100Å, Thermo Scientific) regulated to a temperature of 50°C with a linear gradient of 2% to 30% buffer B (100% MeCN in 0.1% formic acid) at a flow rate of 300 nL/min over 91 min. MS full scans were performed in the ultrahigh-field Orbitrap mass analyzer in ranges m/z 375-1500 with a resolution of 120 000 at m/z 200. The top 20 intense ions were subjected to Orbitrap for further fragmentation via high energy collision dissociation (HCD) activation and a resolution of 15 000 with the AGC target set to 10^5^ ions. We selected ions with charge state from 2+ to 6+ for screening. Normalized collision energy (NCE) was set at 27 and the dynamic exclusion of 40s.

For the Cx30 pull down, LC was performed as previously with an RSLCnano system (same trap column, column and buffers), coupled online to an Orbitrap Exploris 480 mass spectrometer (Thermo Scientific). Peptides were trapped onto a C18 column with buffer A at a flow rate of 3.0 μL/min over 4 min. Separation was performed at a temperature of 40°C with a linear gradient of 3% to 29% buffer B at a flow rate of 300 nL/min over 91 min. MS full scans were performed in the ultrahigh-field Orbitrap mass analyzer in ranges *m*/*z* 375-1500 with a resolution of 120 000 at *m*/*z* 200. The top 20 most intense ions were subjected to Orbitrap for further fragmentation via high energy collision dissociation (HCD) activation and a resolution of 15 000 with the AGC target set to 100%. We selected ions with charge state from 2+ to 6+ for screening. Normalized collision energy (NCE) was set at 30 and the dynamic exclusion of 40s.

##### Data analysis

For identification, the data were searched against the Mus Musculus Swiss - Prot database (UP 000000589 containing 17038 sequences) and also a databank of the common contaminants containing 245 sequences for the Cx30 pull downs using Sequest–HT through proteome discoverer (version 2.2). Enzyme specificity was set to trypsin and a maximum of two-missed cleavage sites were allowed. Oxidized methionine and N-terminal acetylation were set as variable modifications. Maximum allowed mass deviation was set to 10 ppm for monoisotopic precursor ions and 0.02 Da for MS/MS peaks. The resulting files were further processed using myProMS^9^ v3.5 (https://github.com/bioinfo-pf-curie/myproms). FDR calculation used Percolator^10^ and was set to 1% at the peptide level for the whole study. The label free quantification was performed by peptide Extracted Ion Chromatograms (XICs) computed with MassChroQ version 2.2.1^11^. For protein quantification, XICs from proteotypic peptides shared between compared conditions (TopN matching) and missed cleavages of peptides were used. Median and scale normalization was applied on the total signal to correct the XICs for each biological replicate. To estimate the significance of the change in protein abundance, a linear model (adjusted on peptides and biological replicates) based on two-tailed T-tests was performed and P-values were adjusted with a Benjamini-Hochberg FDR. Proteins from the biotinylated WFA lectine pull down with at least three total peptides in all replicates (n=3), a 1.5-fold enrichment and an adjusted P-value < 0.05 were considered significantly enriched in sample comparison. Unique proteins were considered with at least three total peptides in all replicates. Protein selected were further analyzed with Reactome pathway analysis (https://reactome.org/PathwayBrowser/#TOOL=AT,^12^). Considering the lack of functional annotation in Mus Musculus, protein pathways were retrieved from the mapping of selected proteins to Homo Sapiens.

#### Electroretinography

The changes in potentials induced by light flashes in the retinal tissue were recorded with a Siem Medical ERG system (Nîmes, France). The active electrode (a gold loop) was placed on the cornea, while reference and ground electrodes were placed subcutaneously above the skull and in the tail, respectively. Mice were dark-adapted prior to the experiment, being housed in a dark-room on the evening prior to the recordings. On the following day, mice were handled under dim red light to maintain dark adaptation. After pupil dilatation with a drop of atropine and anesthesia with an intraperitoneal injection of a mixture of ketamine + medetomidine hydrochloride, the mice were placed on a mobile platform with a heating pad, to maintain body temperature. Ophthalmic gel was applied on both eyes, to keep the cornea moist and favor electric contact with the active electrode. After placement of the electrodes, the platform was moved forward so that the head of the mice was inside of the Ganzfeld delivering the flashes. After a series of flashes of increasing intensity (from 20 mCd.s.m-2 to 10 Cd. s.m-2), a background light of 25 Cd.m-2 was applied for 15 minutes, to saturate the rod photoreceptors and light adapt the cones, and flashes of 1 and 10 Cd.s.m-2 were applied. The cone system was further tested using flicker stimuli delivered at 2, 10, 15 and 20 Hz, with the same 25 Cd.m-2 background light.

#### Monocular deprivation

For monocular deprivation, mice were anesthetized i.p. with a mix of ketamine (95mg/kg) and xylazine (12mg/kg) in 0.9 % NaCl. The area surrounding the eye to be sutured was trimmed and wiped with 70% ethanol. Two mattress sutures were then placed and animals were checked daily to ensure sutures remained intact.

#### Intrinsic optical imaging

##### Surgery

Mice were anesthetized with urethane (1.2 g/kg, i.p.) and sedated with chlorprothixene (8 mg/kg, i.m.). Atropine (0.1 mg/kg) and dexamethasone (2 mg/kg) were injected subcutaneously. The animals were placed in a stereotaxic apparatus and the body temperature was maintained at 37°C. In some cases, a craniotomy was made over the visual cortex. The exposed area was then covered by agarose (2.5%) and a glass coverslip. In other cases, the skull was left intact and intrinsic signals were recorded directly though the skull. No difference was found between the two methods.

##### Visual stimulation and optical imaging recording

Visual cortical responses were recorded using imaging methods based on Fourier transform following periodic stimulation^13,14^. Visual stimuli were presented on a high refresh rate monitor placed 20 cm in front of the mouse. Stimulation consisted of a horizontal bar drifting downwards periodically at a period of 8 s in the binocular visual field of the recorded hemisphere (from +5° ipsilateral to +15° contralateral). Each eye was stimulated 5 times in alternance for 2 to 4 minutes. Intrinsic signals were recorded using a 1M60 CCD camera (Dalsa) with a 135×50 mm tandem lens (Nikon) configuration. After acquisition of the surface vascular pattern, the camera was focused 400 μm deeper. Intrinsic signals were acquired with a 700 nm illumination wavelength and frames stored at a rate of 10 Hz, after a 2×2 pixels spatial binning.

##### Data analysis

Retinotopic maps for each eye were computed offline. Prior to Fourier transform, the generalized indicator function^15^ was applied to remove slow varying components independent of the stimulation. Retinotopic maps were calculated from the phase and magnitude components of the signal at the frequency of stimulation. For each eye, the five activity maps were averaged, filtered with a Gaussian kernel low-pass filter (3 pixels s.d.) and set with a cut-off threshold at 50% of the maximum response. This allowed defining the binocular zone as the intersection between the response regions of each eye. For each of the 5 repetitions, an OD value at each pixel in the binocular zone was then defined as (C-I)/(C+I), calculated from the response amplitude from the contralateral (C) eye and the ipsilateral (I) eyes. OD values ranged from −1 to 1, with negative values representing an ipsilateral bias, and positive values a contralateral bias. From these 5 OD maps, the OD index was then computed as the average of the OD values in the binocular zone. The OD indexes of one animal were averaged for statistical comparisons between ages, strains and conditions.

#### Slices preparation and electrophysiology

After rapid extraction of the brain, coronal slices (400 μm) containing the visual cortex were cut by mean of a vibratome (Leica VT1200S) and allowed to recover for a minimum of 45 min at 30°C in a submerged chamber containing ACSF before recording. Slices were individually transferred in a recording chamber placed under a microscope (Slicescope SSPro1000) and continuously perfused (2 mL/min) with aCSF composed of (in mM): 119 NaCl, 2.5 KCl, 2.5 CaCl_2_, 1.3 MgSO_4_, 1 NaH_2_PO_4_, 26.2 NaHCO_3_ and 11 glucose. For recording of spontaneous synaptic activity, layer 4 pyramidal neurons from the primary visual cortex were recorded by whole-cell patch clamp using 3-6 MΩ glass pipettes filled with (in mM): 144 potassium gluconate, 1 MgCl_2_,10 HEPES, 0.5 EGTA (pH 7.4, 280 mOsm). Spontaneous excitatory or inhibitory postsynaptic potentials (sEPSCs or sIPSCs) were identified as inward and ourward currents, respectively. Excitation-inhibition (E-I) balance was determined as described previously^16,17^. Briefly, electrical stimulations (10-100 μA, 0.2 ms, 0.05 Hz) were applied in layer 2 of the visual cortex using a 1 MΩ monopolar glass pipette filled with aCSF. Under voltage-clamp, 5 trials were repeated for 5 holding potentials (−70mV to −50mV). Data were analysed off-line with Elphy (NeuroPSI-CNRS, Gif-sur-Yvette, France). The E-I balance determination is based on the continuous measurement of conductance dynamics during the full-time course of the stimulus-evoked synaptic response. Briefly, we performed post-hoc decomposition of postsynaptic current waveforms in excitatory and inhibitory conductances together with continuous estimation of the apparent reversal potential of the composite responses. This allows a somatic measurement of the E-I balance after dendritic integration of incoming excitation and inhibition^17^. In order to extract the excitatory and inhibitory conductance changes from the evoked synaptic currents, the neuron is considered as the point-conductance model of a single-compartment cell, described by the following general membrane equation:

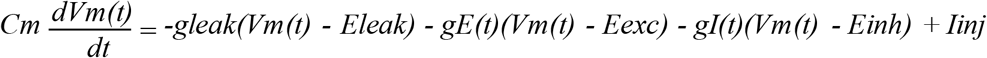

where C_m_ denotes the membrane capacitance, I_inj_ the injected current, g_leak_ the leak conductance and E_leak_ the leak reversal potential. gE(t) and gI(t) are the excitatory and inhibitory conductances, with respective reversal potentials E_exc_ and E_inh_. Evoked synaptic currents were measured and averaged for 5 holding potentials. IV curves were then calculated at all time points of the response. In IV curves for every possible delay (t), the value of holding potential (V_h_) was corrected (V_hc_) from the ohmic drop due to leakage current through the access resistance (V_hc_(t) = V_h_(t) – I(t) x R_s_). An average estimate of the input conductance waveform of the cell was calculated from the best linear fit (mean least square criterion) of the IV curve for each delay (t) following the stimulation onset. Only cells showing a Pearson correlation coefficient for the IV linear regression higher than 0.95 between −90 and −40 mV were considered for calculation of the conductance change in the recorded pyramidal neuron, using the slope of the regression line. The synaptically-evoked global conductance term (g_T_(t)) was then measured by subtracting the resting conductance observed in the absence of stimulation (on a time window of 100 ms before electrical stimulation) from the input total conductance. The synaptic reversal potential of the synaptic conductance (E_syn_(t)) was taken as the voltage of the intersection between the IV curve during the synaptic response and the IV curve at rest. Assuming that the evoked somatic conductance change reflects the composite synaptic input reaching the soma, E_syn_(t) characterizes the stimulation-locked dynamics of the balance between excitation and inhibition. The global synaptic conductance (g_T_(t)) was further decomposed into two conductance components (g_E_(t) and g_I_(t)) corresponding to the activation of excitatory and inhibitory synapses respectively, each associated with known and fixed reversal potentials: 0 mV for excitatory (E_exc_) and −80 mV for inhibitory conductance (E_inh_). Under our experimental conditions, E_syn_(t) took any intermediate values between E_exc_ (0 mV) and E_inh_ (−80 mV) in such a way that the mathematical conditions of the simplification used to calculate g_I_(t) and g_E_(t) were fulfilled. For each component, excitatory and inhibitory, we calculated the conductance change as the mean averaged over a time window of 200 ms. The contribution of each component was expressed by the ratio of its integral value to that of global conductance change.

#### Fasudil treatment

Fasudil hydrochloride (F4660, LC laboratories, Boston, MA, USA) was prepared by diluting 5 mL of solution stock Fasudil 10X (1g Fasudil hydrochloride for 150 mL H2O) in 50 mL of water. The treatment was given for 4 weeks (from P20 to P50) in drinking water (ad libitum; around 3 mg/day/mouse). Drinking bottles were wrapped in aluminium to avoid phototoxicity and replaced every 2 days^18^.

#### Statistical analysis

All data are presented as mean ± standard error of the mean (SEM) and n represent the number of independent replicates. All statistical analyses were performed with GraphPad Prism (GraphPad sofware, USA). Prior statistical comparison, the normality of the distributions were studied and appropriate parametric or nonparametric tests were used. Stastistical significance between two groups was evaluated with a two-tailed t-test or Mann-Whitney test. Concerning within-group comparisons, statistical significance was determined with an ANOVA (One- or two-way) or a Kruskal-Wallis followed by post hoc tests (Tukey or Sidak post-hoc tests for the ANOVA; Dunn post-hoc test for Kruskal-Wallis). Appropriate sample sizes were based on best practices in the literature for each method as well as on ethical standards to minimize numbers of animals for experiments.

## Data availability

We confirm that all relevant data are included in the paper and/or its supplementary information files. The mass spectrometry proteomics data have been deposited to the ProteomeXchange Consortium via the PRIDE^19^ partner repository with the dataset identifier PXD020448 (Username, Password: nmA5W4eW).

## References

1 Espinosa, J. S. & Stryker, M. P. Development and plasticity of the primary visual cortex. Neuron 75, 230–249, doi:10.1016/j.neuron.2012.06.009 (2012).

2 Hubel, D. H. & Wiesel, T. N. The period of susceptibility to the physiological effects of unilateral eye closure in kittens. J Physiol 206, 419–436, doi:10.1113/jphysiol.1970.sp009022 (1970).

3 Hensch, T. K. Critical period regulation. Annu Rev Neurosci 27, 549–579, doi:10.1146/annurev.neuro.27.070203.144327 (2004).

4 LeBlanc, J. J. & Fagiolini, M. Autism: a “critical period” disorder? Neural Plast 2011, 921680, doi:10.1155/2011/921680 (2011).

5 Meredith, R. M. Sensitive and critical periods during neurotypical and aberrant neurodevelopment: a framework for neurodevelopmental disorders. Neurosci Biobehav Rev 50, 180–188, doi:10.1016/j.neubiorev.2014.12.001 (2015).

6 Doll, C. A. & Broadie, K. Impaired activity-dependent neural circuit assembly and refinement in autism spectrum disorder genetic models. Front Cell Neurosci 8, 30, doi:10.3389/fncel.2014.00030 (2014).

7 Clarke, L. E. & Barres, B. A. Emerging roles of astrocytes in neural circuit development. Nat Rev Neurosci 14, 311–321, doi:10.1038/nrn3484 (2013).

8 Lehmann, K. & Lowel, S. Age-dependent ocular dominance plasticity in adult mice. PLoS One 3, e3120, doi:10.1371/journal.pone.0003120 (2008).

9 Muller, C. M. & Best, J. Ocular dominance plasticity in adult cat visual cortex after transplantation of cultured astrocytes. Nature 342, 427–430, doi:10.1038/342427a0 (1989).

10 Cahoy, J. D. et al. A transcriptome database for astrocytes, neurons, and oligodendrocytes: a new resource for understanding brain development and function. J Neurosci 28, 264–278, doi:10.1523/JNEUROSCI.4178-07.2008 (2008).

11 Ghezali, G. et al. Connexin 30 controls astroglial polarization during postnatal brain development. Development 145, doi:10.1242/dev.155275 (2018).

12 Pizzorusso, T. et al. Reactivation of ocular dominance plasticity in the adult visual cortex. Science 298, 1248–1251, doi:10.1126/science.1072699 (2002).

13 Erchova, I., Vasalauskaite, A., Longo, V. & Sengpiel, F. Enhancement of visual cortex plasticity by dark exposure. Philos Trans R Soc Lond B Biol Sci 372, doi:10.1098/rstb.2016.0159 (2017).

14 He, H. Y., Ray, B., Dennis, K. & Quinlan, E. M. Experience-dependent recovery of vision following chronic deprivation amblyopia. Nat Neurosci 10, 1134–1136, doi:10.1038/nn1965 (2007).

15 Hofer, S. B., Mrsic-Flogel, T. D., Bonhoeffer, T. & Hubener, M. Lifelong learning: ocular dominance plasticity in mouse visual cortex. Curr Opin Neurobiol 16, 451–459, doi:10.1016/j.conb.2006.06.007 (2006).

16 Di Cristo, G. et al. Activity-dependent PSA expression regulates inhibitory maturation and onset of critical period plasticity. Nat Neurosci 10, 1569–1577, doi:10.1038/nn2008 (2007).

17 Hensch, T. K. et al. Local GABA circuit control of experience-dependent plasticity in developing visual cortex. Science 282, 1504–1508, doi:10.1126/science.282.5393.1504 (1998).

18 Tropea, D. et al. Gene expression changes and molecular pathways mediating activity-dependent plasticity in visual cortex. Nat Neurosci 9, 660–668, doi:10.1038/nn1689 (2006).

19 Turrigiano, G. G. & Nelson, S. B. Homeostatic plasticity in the developing nervous system. Nat Rev Neurosci 5, 97–107, doi:10.1038/nrn1327 (2004).

20 Gonzalez-Billault, C. et al. The role of small GTPases in neuronal morphogenesis and polarity. Cytoskeleton (Hoboken) 69, 464–485, doi:10.1002/cm.21034 (2012).

21 Kim, H. J. et al. RhoA/ROCK Regulates Prion Pathogenesis by Controlling Connexin 43 Activity. Int J Mol Sci 21, doi:10.3390/ijms21041255 (2020).

22 Mendoza-Naranjo, A. et al. Targeting Cx43 and N-cadherin, which are abnormally upregulated in venous leg ulcers, influences migration, adhesion and activation of Rho GTPases. PLoS One 7, e37374, doi:10.1371/journal.pone.0037374 (2012).

23 Jeong, K. J. et al. The Rho/ROCK pathway for lysophosphatidic acid-induced proteolytic enzyme expression and ovarian cancer cell invasion. Oncogene 31, 4279–4289, doi:10.1038/onc.2011.595 (2012).

24 Tong, L. & Tergaonkar, V. Rho protein GTPases and their interactions with NFkappaB: crossroads of inflammation and matrix biology. Biosci Rep 34, doi:10.1042/BSR20140021 (2014).

25 Pannasch, U. et al. Connexin 30 sets synaptic strength by controlling astroglial synapse invasion. Nat Neurosci 17, 549–558, doi:10.1038/nn.3662 (2014).

## References

1 Tsai, H. H. et al. Regional astrocyte allocation regulates CNS synaptogenesis and repair. Science 337, 358–362, doi:10.1126/science.1222381 (2012).

2 Boulay, A. C. et al. Hearing is normal without connexin30. J Neurosci 33, 430–434, doi:10.1523/JNEUROSCI.4240-12.2013 (2013).

3 Zhuo, L. et al. hGFAP-cre transgenic mice for manipulation of glial and neuronal function in vivo. Genesis 31, 85–94, doi:10.1002/gene.10008 (2001).

4 Rouach, N., Calvo, C. F., Glowinski, J. & Giaume, C. Brain macrophages inhibit gap junctional communication and downregulate connexin 43 expression in cultured astrocytes. Eur J Neurosci 15, 403–407, doi:10.1046/j.0953-816x.2001.01868.x (2002).

5 Qamar Saeed, M. et al. Comparison Between Several Integrase-defective Lentiviral Vectors Reveals Increased Integration of an HIV Vector Bearing a D167H Mutant. Mol Ther Nucleic Acids 3, e213, doi:10.1038/mtna.2014.65 (2014).

6 Berger, A. et al. Repair of rhodopsin mRNA by spliceosome-mediated RNA trans-splicing: a new approach for autosomal dominant retinitis pigmentosa. Mol Ther 23, 918–930, doi:10.1038/mt.2015.11 (2015).

7 Cahoy, J. D. et al. A transcriptome database for astrocytes, neurons, and oligodendrocytes: a new resource for understanding brain development and function. J Neurosci 28, 264–278, doi:10.1523/JNEUROSCI.4178-07.2008 (2008).

8 Pillet, L. E. et al. The intellectual disability protein Oligophrenin-1 controls astrocyte morphology and migration. Glia 68, 1729–1742, doi:10.1002/glia.23801 (2020).

9 Poullet, P., Carpentier, S. & Barillot, E. myProMS, a web server for management and validation of mass spectrometry-based proteomic data. Proteomics 7, 2553–2556, doi:10.1002/pmic.200600784 (2007).

10 The, M., MacCoss, M. J., Noble, W. S. & Kall, L. Fast and Accurate Protein False Discovery Rates on Large-Scale Proteomics Data Sets with Percolator 3.0. J Am Soc Mass Spectrom 27, 1719–1727, doi:10.1007/s13361-016-1460-7 (2016).

11 Valot, B., Langella, O., Nano, E. & Zivy, M. MassChroQ: a versatile tool for mass spectrometry quantification. Proteomics 11, 3572–3577, doi:10.1002/pmic.201100120 (2011).

12 Jassal, B. et al. The reactome pathway knowledgebase. Nucleic Acids Res 48, D498–D503,, doi:10.1093/nar/gkz1031 (2020).

13 Cang, J. et al. Development of precise maps in visual cortex requires patterned spontaneous activity in the retina. Neuron 48, 797–809, doi:10.1016/j.neuron.2005.09.015 (2005).

14 Kalatsky, V. A. & Stryker, M. P. New paradigm for optical imaging: temporally encoded maps of intrinsic signal. Neuron 38, 529–545, doi:10.1016/s0896-6273(03)00286-1 (2003).

15 Yokoo, T., Knight, B. W. & Sirovich, L. An optimization approach to signal extraction from noisy multivariate data. Neuroimage 14, 1309–1326, doi:10.1006/nimg.2001.0950 (2001).

16 Meunier, C. N. et al. D-Serine and Glycine Differentially Control Neurotransmission during Visual Cortex Critical Period. PLoS One 11, e0151233, doi:10.1371/journal.pone.0151233 (2016).

17 Monier, C., Fournier, J. & Fregnac, Y. In vitro and in vivo measures of evoked excitatory and inhibitory conductance dynamics in sensory cortices. J Neurosci Methods 169, 323–365, doi:10.1016/j.jneumeth.2007.11.008 (2008).

18 Meziane, H. et al. Fasudil treatment in adult reverses behavioural changes and brain ventricular enlargement in Oligophrenin-1 mouse model of intellectual disability. Hum Mol Genet 25, 2314–2323, doi:10.1093/hmg/ddw102 (2016).

19 Perez-Riverol, Y. et al. The PRIDE database and related tools and resources in 2019: improving support for quantification data. Nucleic Acids Res 47, D442–D450,, doi:10.1093/nar/gky1106 (2019).

